# The swine spatiotemporal H3K27ac spectrum provides novel resources for exploring gene regulation related to complex traits and fundamental biological process

**DOI:** 10.1101/2021.07.28.454245

**Authors:** Yaling Zhu, Zhimin Zhou, Tao Huang, Zhen Zhang, Wanbo Li, Ziqi Ling, Tao Jiang, Jiawen Yang, Siyu Yang, Yanyuan Xiao, Carole Charlier, Michel Georges, Bin Yang, Lusheng Huang

## Abstract

The limited knowledge of genomic non-coding and regulatory regions has limited our ability to decipher the genetic mechanisms underlying complex traits in pigs. In this study, we characterize the spatiotemporal landscape of putative enhancers and promoters and their target genes by combining H3K27ac targeted ChIP-Seq and RNA-Seq in fetal (day 74-75 pc) and adult (day 132-150 pn) tissues (brain, liver, heart, muscle and small intestine) sampled from Asian aboriginal Bamaxiang and European highly selected Large White pigs of both sexes. We identify 101,290 H3K27ac peaks marking 18,521 promoters and 82,769 enhancers, including peaks that are active across all tissues and developmental stages could indicate safe harbors for exogenous gene insertion, and tissue and developmental-stage specific peaks that regulate genes pathways matching tissue and developmental stage specific physiological functions. We found H3K27ac and DNA methylation in the promoter region of the *XIST* gene may involve in X chromosome inactivation, and demonstrate utility of the present resource to reveal regulatory patterns of known causal genes and to prioritize candidate causal variants for complex traits in pigs. We have developed a web browser to improve the accessibility of the results (http://39.108.231.116/browser/?genome=susScr11).

## Introduction

It is increasingly apparent that a substantial proportion of the heritability for quantitative traits, including economically important traits in livestock (Georges et al., 2019), is attributable to variants affecting gene regulatory elements including proximal promoters, distant enhancers and silencers. These regulatory elements have been extensively explored in multiple tissues/cell types at different developmental stages in human (Roadmap Epigenomics et al., 2015) and mice (Mouse et al., 2012) through approaches targeting whole genome open chromatin (Dnase I and ATAC-Seq) and histone modifications (ChIP-Seq) followed by next generation sequencing. These data have contributed to a better understanding of gene regulation underlying tissue or cell lineage differentiation (Roadmap Epigenomics, et al., 2015) and development (Nord et al., 2013). Moreover, intersecting the landscape of regulatory regions with risk loci identified by genome wide association studies (GWAS) revealed tissues and cell types relevant to disease traits (Roadmap Epigenomics, et al., 2015), as well as regulatory mechanisms underlying disease associated loci with large effects (Nasrallah et al., 2020).

A recent study has shown that ubiquitous chromatin opening elements (UCOE) marked by e.g. Histone 3 lysine 27 acetylation (H3K27ac) and DNase I hypersensitivity marks can support stable expression of nearby genes (Neville et al., 2017). This makes UCOE informative for the identification of safe harbors, preferably in intergenic regions, in which exogenous genes can be inserted and stably expressed across tissues and development stages. Identifying putative safe harbors would be very helpful for the production of transgenic animals for manufacturing therapeutic proteins and for other biomedical studies (Neville, et al., 2017). Currently, the major safe harbors used in swine models are Rosa26 (Li et al., 2014) and H11 (Ruan et al., 2015), which were first identified in the mouse (Friedrich and Soriano, 1991, Tasic et al., 2011). No study has been implemented to scan the swine genome for candidate safe harbors based on chromatin modification and gene expression data in the pig.

Part of the mammalian regulatory elements reside within the ∼5% of the genome under evolutionary constraint (Lindblad-Toh et al., 2011). Thus, inter-species sequence conservation effectively contributes to the identification of at least a portion of mammalian regulatory space. However, distant regulatory elements in particular have been shown to be capable of rapid evolutionary turnover despite their functional importance (Nord, et al., 2013, Villar et al., 2015). Therefore, it is essential to identify gene regulatory elements at genome-wide scale independent of inter-species conservation. Towards that goal, collaborative initiatives are now underway in livestock species, collectively referred to as the Functional Annotation of ANimal Genome (FAANG) project (Andersson et al.). In pigs, studies have profiled H3K27ac and H3K4me3 in bulk liver tissue (Villar, et al., 2015), placental tissues of gestational days 50 and 95 fetuses(Han et al., 2019), and open chromatin regions (ATAC-Seq) and genome 3D topology (Hi-C) in CD4+ and CD8+ T cells, and dissociated liver cells(Foissac et al., 2019). A recent study performed H3K27ac and H3K4me4 ChIP-Seq on 12 tissues, as well as ATAC-seq on muscle and fat tissues from four pig breeds, revealing more than 220,000 cis-regulatory elements in the pig genome (Zhao et al., 2021). Yet, further efforts are needed to enrich the catalogue of pig regulatory elements in multiple tissues at different developmental stages.

We herein describe the characterization of H3K27ac profiles across the porcine genome. H3K27ac is an established histone mark of active promoters and enhancers, which was showed to be highly variable across tissues and significantly enriched for disease associated variants in humans(Roadmap Epigenomics, et al., 2015). Assays were performed on 70 samples corresponding to five tissues (brain, heart, liver, muscle and small intestine), two developmental stages (fetal 74-75 days post conceptus and adult 132-150 days postnatal), two breeds (European Large White and Chinese Bama Xiang) and the two sexes of pigs (**Fig. 1A**). Sixty-two of these samples were also profiled by RNA sequencing. We obtained a total of 101,290 H3K27ac peaks evidenced in at least two samples that marked 18,521 active promoters and 82,769 active enhancers. These peaks were further characterized for their 1) genomic distribution, 2) association with expression of nearby genes, 3) constitutive activity across different tissue - developmental stages, and their utilization in revealing potential safe harbors for genome editing, 4) specific activities in different tissue and developmental stages, and 5) differential activity between the two breeds and two sexes. We also demonstrate the utility of H3K27ac peaks to identify regulatory patterns of causal genes and causative variants underlying complex traits.

**Fig. 1:**
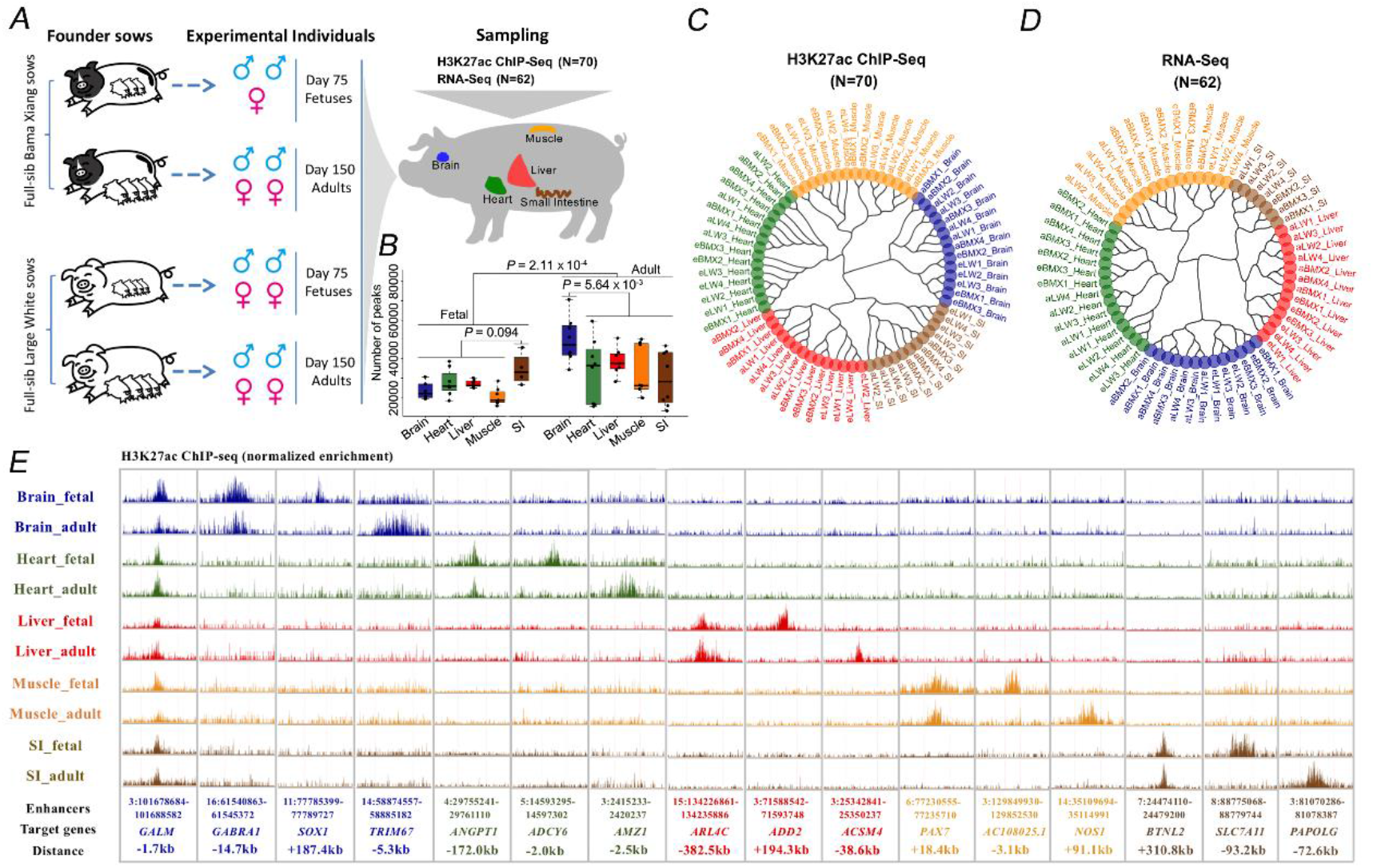
Identifying the activity landscape of active promoters and enhancers via H3K27ac ChIP-Seq in pigs. (**A)** Schematic diagram showing the design, samples and data collected in this study. (**B)** Boxplot showing the distribution of the number of peaks identified in samples sorted by tissue type and developmental stage. (**C-D)** Neighbor joining tree of samples inferred from peak activity and gene expression seperates the ten tissues x developemental stage groups (5 tissues × 2 developmental stages). (**E)** Representative examples of H3K27ac peaks showing (i) constitutive activity: near *GALM*, a gene encodes an enzyme involved in the fundamental cellular function of galactose metabolism(Vaillancourt et al., 2008), and 5 tissue specific activity with or without additional developmental stage specific activity: brain specific peaks locate close to *GABRA1* (neurotransmitter in mammalian brain), *SOX1* (embryonic and forebrain neuron development) and *TRIM67* (Brain development and behavior) with function matching the biology of brain; heart specific peaks locate close to *ANGPT1I* (angiopoietin and heart development), *ADCY6* (adenylyl cyclase related to vascular sensitivity) and *AMZ1* (heart development); liver specific peaks near *ARL4C* (cholesterol transport), *ADD2* (hematopoietic protein) and *ACSM4* (fatty acid ligase); muscle specific peaks near *PAX7* (transcription factor regulating myogenesis), *AC1080251* (a transcript with unknown function) and *NOS1* (Skeletal muscle integrity and contractile); small intestine specific peaks near *BTNL2* (immune surveillance and ulcerative colitis), *SLC7A11* (amino acid transmembrane transport) and *PAPOLG* (immune process).

## Results

### The H3K27ac landscape in five tissues, two sexes, two breeds and two developmental stages of the pig

Two Large White full-sisters (sow1 and sow2) were mated to the same Large White boar. Sow 1 was slaughtered at day 75 post mating, and two male and two female fetuses were harvested. Sow2 was raised until delivery and two male and two female offspring slaughtered at 150 days after birth for sampling. The same design was also applied to the Chinese aboriginal Bama Xiang pig breed, except that only 3 fetuses were harvested at day 74 and that the adult Bama Xiang pigs were slaughtered at day 132-141. We sampled brain, liver, heart, muscle and small intestine from all fetuses and adult animals (**Fig. 1A**).

We performed H3K27ac ChIP-Seq on 70/75 samples and RNA-Seq on 62/75 samples (**Fig. 1A and Supplementary Table 1&2)**. For the ChIP-Seq data, an average of 31.2 and 30.3 million single-end 50-bp reads were obtained for input and immune-precipitated (IP) samples, respectively, of which 87.5% and 89.5% were mapped to the Sus scrofa genome assembly 11.1 **(Supplementary Table 1 and Methods)**. From these data, we identified an average of 31,121 peaks per sample (range: 13,064 to 70649), with average peak length of 2,527 bp (range: 1,224 to 5,480 bp), which is comparable to previous reports(Villar, et al., 2015) **(Supplementary Table 1 and Supplementary Fig. 1)**. Adult samples have significantly higher peak numbers than fetal samples (*P* = 2.1×10^-4^) **(Fig. 1B)**. Moreover, the adult brain samples have more peaks on average than the other adult tissues (*P* = 5.6×10^-3^) **(Fig. 1B)**, which may reflect the larger cellular heterogeneity of the brain compared to other tissues.

For the RNA-Seq data, an average of 43.3 million reads were generated for the 62 samples, of which an average of 93.8% mapped to the reference genome **(Supplementary Table 2)**. Transcripts were assigned to a reference set of 44,415 genes (34,355 from Ensembl and 10,060 from de novo assembled by Stringtie (**Supplementary Table 3**). After quality control procedures (Materials and Methods), 20,319 genes were retained for further analysis. Gene expression levels were normalized using the VST approach in DESeq2 (Love et al., 2014).

To facilitate comparison of H3K27ac across samples, we combined peaks across samples and removed the largest 5% in terms of peak length to yield a set of 170,371 merged peaks, which have an average length of 2,667 bp, only marginally larger than the average length of peaks (2,527 bp) identified in individual samples before merging. First, we used a binary way to define the “active” and “inactive” statuses of a peak, and considered that a merged peak is active in a given sample if it overlaps by at least 1 base pair with any peak identified in that sample. We further measured the activities of merged peaks in each sample quantitatively as the average read depths per million IP reads minus average read depths per million input reads in the peak region. It is noteworthy that the binarily inactive peaks in an individual could be covered by sequence reads, and that therefore their quantitative activity could be greater than zero. Based on the binary peak activity, 85.3% of peaks active in at least one of our adult male liver samples overlapped with the adult male liver peaks identified in an independent study (Villar, et al., 2015) (**Supplementary Fig. 2**), supporting the validity of the peaks revealed in this study. The clustering analysis based on the quantitative activities of the 170,371 peaks showed that the samples first grouped by tissue, followed by developmental stage without exceptions **(Fig. 1C and Supplementary Fig. 3A)**. A similar clustering pattern was observed based on the expression level of 20,319 genes from RNA-Seq data, except for heart for which breed effect appeared stronger than developmental stage effect **(Fig. 1D and Supplementary Fig. 3B)**. Thus, both H3K27ac and gene expression profiles accurately predict tissue type and developmental stage of the samples. We exemplify peaks that are ubiquitously active across all tissues and developmental stages, as well as peaks that are tissue- but not developmental-stage specific, and peaks that are tissue- and developmental stage-specific (**Fig. 1E**).

We next focused on 101,290 peaks that were active (binary phenotype) in at least any two samples. Following Nord et al.(Nord, et al., 2013), we classified 18,521 peaks intersecting with the 5’ end of any transcript (+/- kb) inferred from the RNA-Seq data or provided by the Ensembl database (release 98) as proximal peaks/putative promoters. The remaining 82,769 were considered as distal peaks/putative enhancers. The quantitative activity of the 18,521 proximal peaks within each sample were significantly correlated with the expression level of their cognate genes (average Pearson correlation coefficient = 0.37, *P* value < 1×10^-300^) (**Supplementary Fig. 4**), reflecting a link between a gene’s expression level and the degree of H3K27 acetylation of its promoter region.

We observed that the distal peaks were mostly located in introns (59.5%), followed by intergenic (34.4%) regions. Among the intronic peaks, more than half are located in the first three introns of the corresponding genes **(Supplementary Fig. 5)**. This underscores the important role of intronic regions close to the 5’ end of genes in regulation of gene transcription, in agreement with reports in other species like rat(Chan et al., 1999) and Arabidopsis(Gallegos and Rose, 2017). Compared to proximal peaks, distal peaks were more likely to be active in a specific tissue in both fetal and adult samples **(Supplementary Fig. 6)**. Similar results were also observed in mice(Nord, et al., 2013), reflecting the important roles of enhancers in controlling tissue specific gene expression. Moreover, distal peaks have significant lower GC content, extent of sequence conservation across species (GERP score), peak length and peak activity compared to proximal peaks, reflecting their distinct biological identity (**Supplementary Fig. 7**).

### Identifying target genes of H3K27ac regions across the whole genome and potential silencing role of H3K27ac on gene transcription

To investigate the associations of H3K27ac elements with nearby gene expression, we calculated the correlations of each H3K27ac peak with expression levels of genes within 500 kb region from corresponding peak across 62 samples, given that 75% of promoter based DNA interaction revealed by HiC data were reported to be within 500 kb (Javierre et al., 2016). We identified an estimated 41,105 correlations, including 35,644 positive and 5,461 negative out of 1089891 tests at *q* value threshold of 0.05. We next investigated the distribution of positive and negative peak-gene correlations against peak-gene distances **(Fig. 2)**. The number of significant correlations for peaks located upstream and downstream of their associated genes appeared to be approximately symmetric, suggesting that the H3K27ac regions can act equally well upstream and downstream of transcript start sites (TSSs) of corresponding genes (**Fig. 2A**). Interestingly, the positive correlations were enriched in peaks and genes located nearby (**Fig. 2A**), while the negative peak-gene correlations appeared to be underrepresented for peaks close to their target genes (**Fig. 2B**), reflecting a mechanism that H3K27ac at promoter regions are always associated with increased expression of corresponding genes. Moreover, the peak length is positively associated with the strength of peak-gene correlations particularly for distal peaks (**Fig. 2C-F**). We reasoned that the peaks with greater length could have extended open chromatin region that better support the transcription of corresponding genes, and thereby contributed to stronger peak – gene correlations. An alternative explanation is that the peak activity is better estimated for large than for short peaks, as more reads are used to estimate the activity of large peaks than for small peaks.

**Fig. 2:**
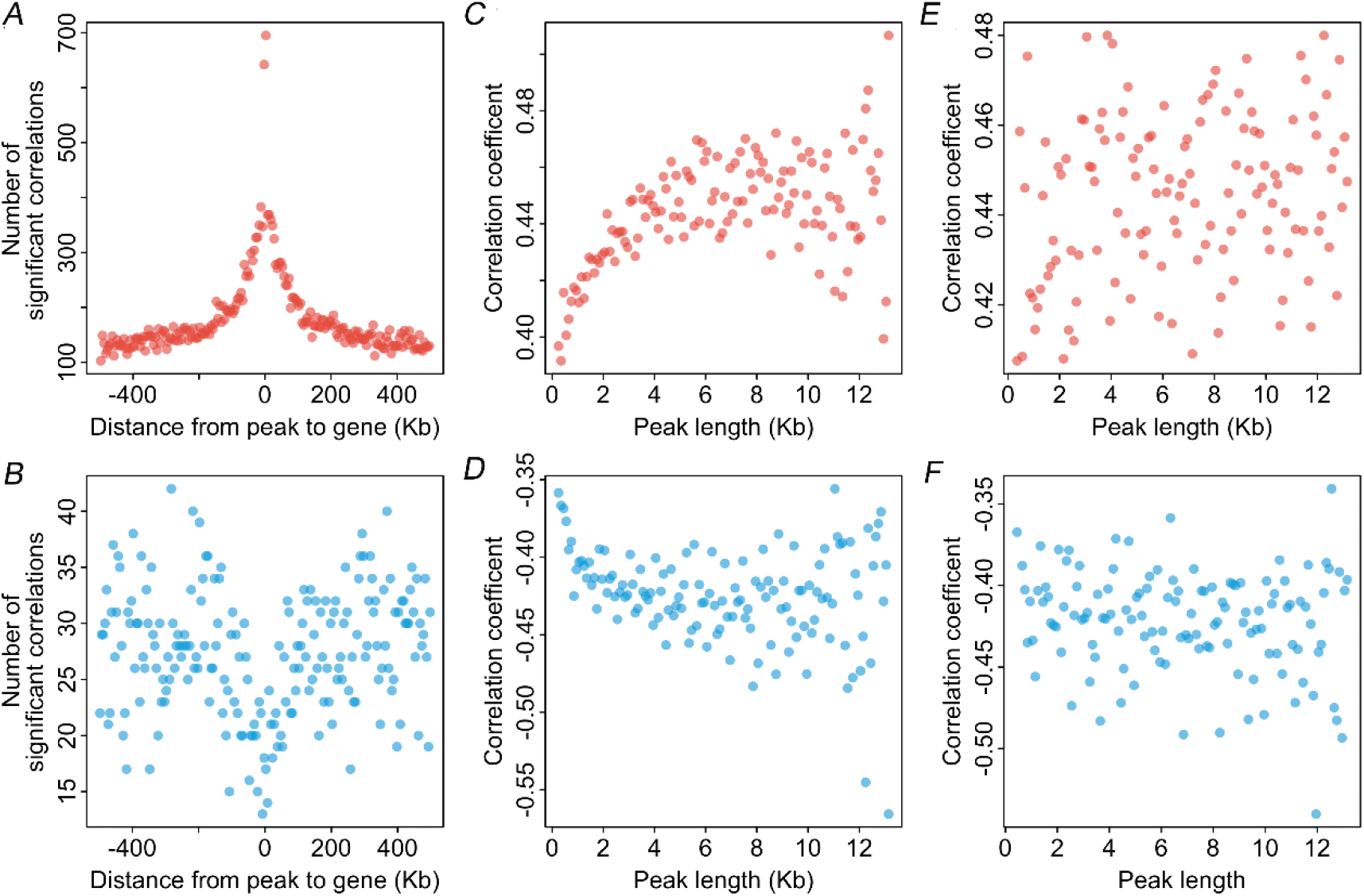
Characterizing the correlations between peak activity and expression of nearby genes (within 500 kb). (**A)** Distribution of significant positive peak-gene correlations as a function of peak-gene distance in a window of 1Mb; (**B)** Distribution of significant negative peak-gene correlations as a function of peak-gene distance; The dots represent number of significant correlations in 5 kb sliding windows. (**C-D)** The distribution of positive and negative correlation coefficients as a function of peak size (distal peaks) in a window of 1Mb. (**E-F)** The distribution of positive and negative correlation coefficients as a function of peak size (proximal peaks); (**C-F)** The results were visualized by 100 bp sliding windows in terms of peak size.

### H3K27ac peaks that are ubiquitously active may serve as safe harbors for swine genome editing

We next searched for peaks that are ubiquitously active across tissues, ages, sex and breeds. We assumed that such peaks may play important roles in maintaining fundamental biological functions, and may highlight safe harbors where transgenes could be inserted and stably expressed. We defined a peak as ubiquitously active if it was active in at least half of samples in each of the 10 tissue x developmental stage groups (**Fig. 1C and methods**). Based on this, we identified a total of 2,679 ubiquitously active peaks, of which most (96.7%) were proximal. Agreeing with our expectations, the genes whose TSS overlapped with ubiquitously active peaks were enriched for basic housekeeping functions such as metabolism of proteins (*q* = 1.2×10^- 58^), RNA metabolism (*q* = 7.1×10^-24^) and splicing (*q* = 1.6×10^-12^) (**Supplementary Table S4 and Fig. 3A**).

**Fig. 3:**
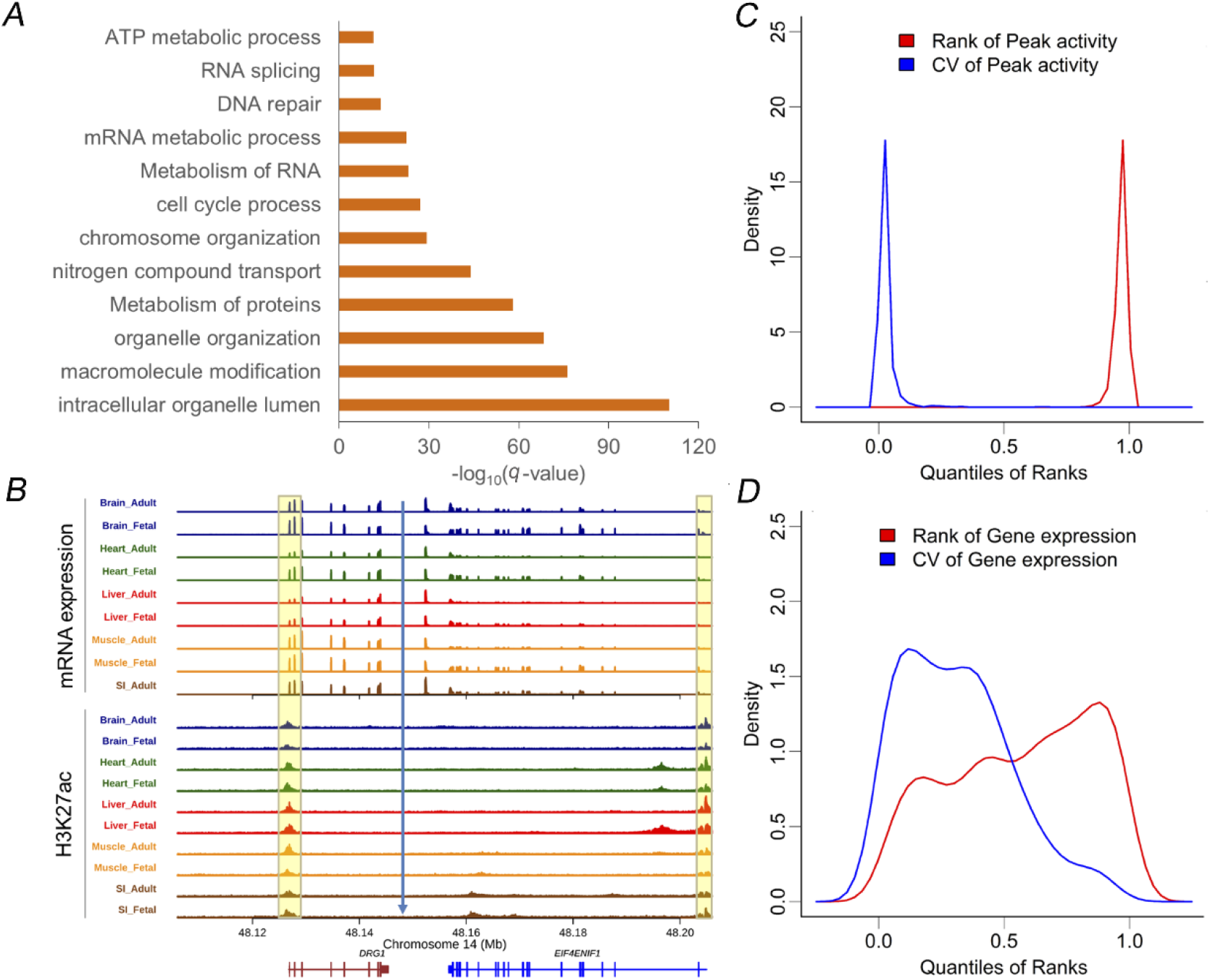
Properties of ubiquitously active H3K27ac peaks and their potential to identify safe harbors. (**A)** Biological processes enriched in cognate genes of the ubiquitously active peaks (the genes whose TSS overlap with the peaks). (**B)** The tracks of mRNA expression and H3K27ac across the tissue – developmental stages at H11 safe harbor locus on pig genome. The transparent yellow bars mark the peaks that show constitutive activity across tissue x developmental stages. The blue arrow marks a potential intergenic safe harbor, a region between two ubiquitously expressed genes, for exogenous gene insertion; (**C)** The distributions of peak activity and coefficient of variations of activity for the 2,679 ubiquitously active peaks defined in this study. (**D)** The distributions of expression levels and coefficient of variation of expression for genes cognate to the 2,679 ubiquitously active peaks.

To test if the ubiquitously active peaks can help to identify potential safe harbors in pigs, we compared expression means and coefficient of variations of genes adjacent to the ubiquitously active peaks with those of all other genes, and found that the former showed significantly higher average expression levels (*P* value < 4.5×10^-13^) and lower CV (*P* value < 2.2×10^-16^) across the 62 samples investigated (**Fig. 3C-D**), suggesting that the ubiquitously active peaks could support high and stable expression of neighboring genes. We further examined the peak activity and gene expression of two known safe harbors (Papapetrou and Schambach, 2016): Rosa26 and H11, and found that both Rosa26 and H11 regions contained peaks and genes that are active or highly expressed across all tissues and development stages (**Supplementary Fig. 8 and Fig. 3B**). Encouraged by these results, we further filtered out 110 ubiquitously active peak regions containing genes with larger means and smaller CV of expression compared to the *THUMPD3* gene in the Rosa26 region and the *DRG1* gene in the H11 region as candidate safe harbors (**Supplementary Table 5**). Two such regions are exemplified in **Supplementary Fig. 9.** Overall, the H3K27ac and RNA-Seq data across tissues, ages, sexes and breeds provides a valuable resource to identify genome wide potential safe harbors for exogenous gene insertion in genome engineering of pigs.

### Tissue specific peaks associate with biological processes and transcription factors matching tissue specific physiological functions

We next investigated the impact of tissue, development stage, breed and sex on peak activity using a linear mixed model. It showed that the tissue, developmental stage, breed and sex on average explained 21.2%, 4.0%, 0.01% and 0.35% of variation in peak activity and 51.1%, 6.1%, 0.94% and 0.02% of variation in mRNA expression, respectively **(Supplementary Fig. 10**). This demonstrates that the tissue is the largest source of variation underlying both peak activity and gene expression, followed by development stage, while the impact of breed and sex is much lower.

To identify H3K27ac peaks that potentially regulate tissue specific gene expression and in turn tissue specific physiological functions, we searched for peaks that are active in one specific tissue in both fetal and adult stages. We selected peaks that were (i) active (binary) in at least half of both fetal and adult samples for the target tissue without being active (binary) in any other sample, and had average activity (quantitative) that was at least four times higher in the target tissue when compared to all other tissues. We identified a total of 790, 420, 940, 156 and 259 tissue specific peaks in brain, heart, liver, muscle and small intestine, respectively (**Fig. 4A**). All were significantly associated with their respective tissue type in a linear regression analyses at *q* value threshold of 0.05, supporting the validity of our filtering approach. Compared to the ubiquitously active peaks, the tissue specific peaks were mainly distal (**Supplementary Fig.11**), reflecting an important role of distal peaks in determining tissue specific biology.

**Fig. 4:**
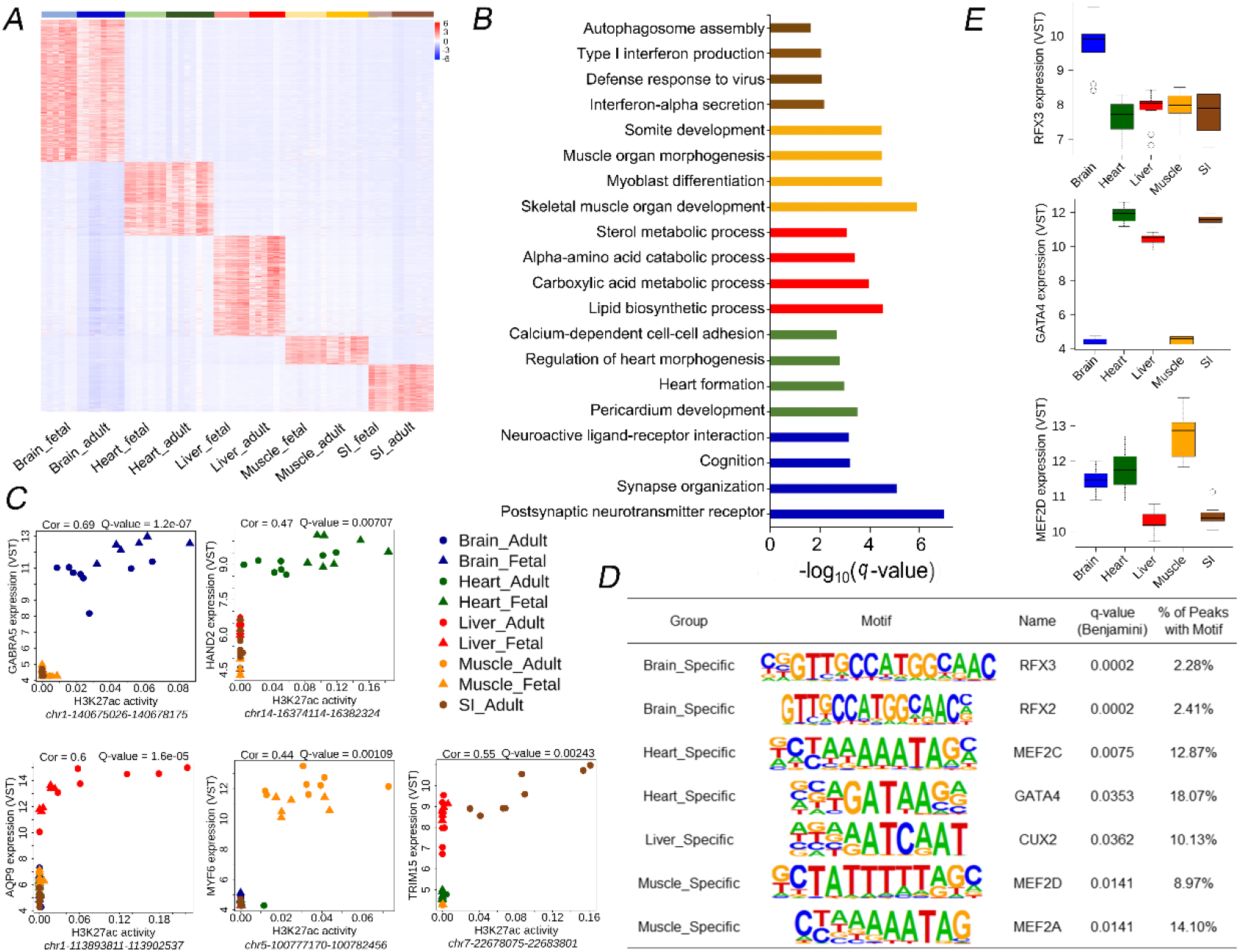
Functional annotation of tissue specific H3K27ac peaks in the five tissues. (**A)** Heatmap of tissue specific peaks in the five tissues. Each row of the heatmap shows the normalized density of read-depth at one peak. (**B)** Representative examples of enriched biological processes in target genes of tissue-specific peaks. (**C)** Representative examples of correlation of tissue specific peak activity with nearby gene expression. (**D)** List of transcription factors with binding motifs enriched in tissue specific peaks. (**E)** Transcription factors whose binding sites are enriched in tissue-specific peaks are highly expressed in the correponding tissue: examples. The blue, green, red, orange and tan colors indicate signals in brain, heart, liver, muscle and small intestine, respectively, the light colors in the lower three graphs indicate fetal stages of corresponding tissues.

The gene ontology enrichment analysis for the target genes of the tissue specific peaks (i.e. the gene within 500Kb from the peak with the highest correlation between peak activity and expression) revealed biological processes such as postsynaptic neurotransmitter receptor activity (*q* = 9.8×10^-8^) for brain-specific peaks, heart formation (*q* = 0.001) for heart-specific peaks, lipid biosynthetic process (*q* = 2.8×10^-5^) and cellular amino acid metabolic process (*q* = 9.9×10^-5^) for liver-specific peaks, skeletal muscle organ development (*q* = 1.2×10^-6^) for muscle-specific peaks, and interferon-alpha secretion (*q* = 6.7×10^-3^) for SI-specific peaks (**Fig. 4B-C, Supplementary Fig. 12 and Supplemental Table 6**). Transcription factors play central roles in tissue differentiation and development(Almalki and Agrawal, 2016). We further performed enrichment analysis of TFBS in the tissue specific peaks, which revealed an overrepresentation of TFs related to the biology of corresponding tissues. These include RFX3 and RFX2 in brain, MEF2C, MEF2D, GATA2, GATA4, MEF2A and ESRRG in heart, CUX2 and NR2f2 in liver, and MEF2D and MEF2A in muscle (**Fig. 4D and Supplementary Table 7**). Among these transcription factors, RFX3 showed highest expression in brain, GATA2, GATA4 and MEF2A showed highest expression in heart, and MEF2D showed highest expression in muscle (**Fig. 4E and Supplementary Fig. 13**). Taken together, we found that tissue specific peaks were associated with genes involved in biological processes and bound by transcription factors related to the biology of corresponding tissues, emphasizing the critical roles of tissue specific H3K27ac peaks in driving tissue specific biological functions.

### Developmental stage specific H3K27ac peaks in the five tissues

We next sought to identify peaks showing developmental stage specific activity, and their potential association with pathways underlying the functional switch from fetal to adult stage (**Fig. 5A**). We defined fetal/adult specific peaks of a tissue as those (i) active (binary) in at least half of fetal/adult samples, but inactive in all adult/fetal samples, and with average quantitative activity at least four-fold different between the two developmental stages of the corresponding tissue. Based on this, we identified 428, 949, 2145, 1202 and 2705 fetal-specific, and 4328, 1376, 5778, 2522 and 973 adult-specific peaks in brain, heart, liver, muscle and duodenum, respectively (**Fig. 5A-B**). The developmental stage specific peaks are mostly tissue specific (95.9% for the fetal and 89.6% for the adult specific peaks) (**Fig. 5B**), as only 1 fetal and 17 adult specific peaks were shared amongst four tissues (**Fig. 5B)**, while none was shared across the five tissues.

**Fig. 5:**
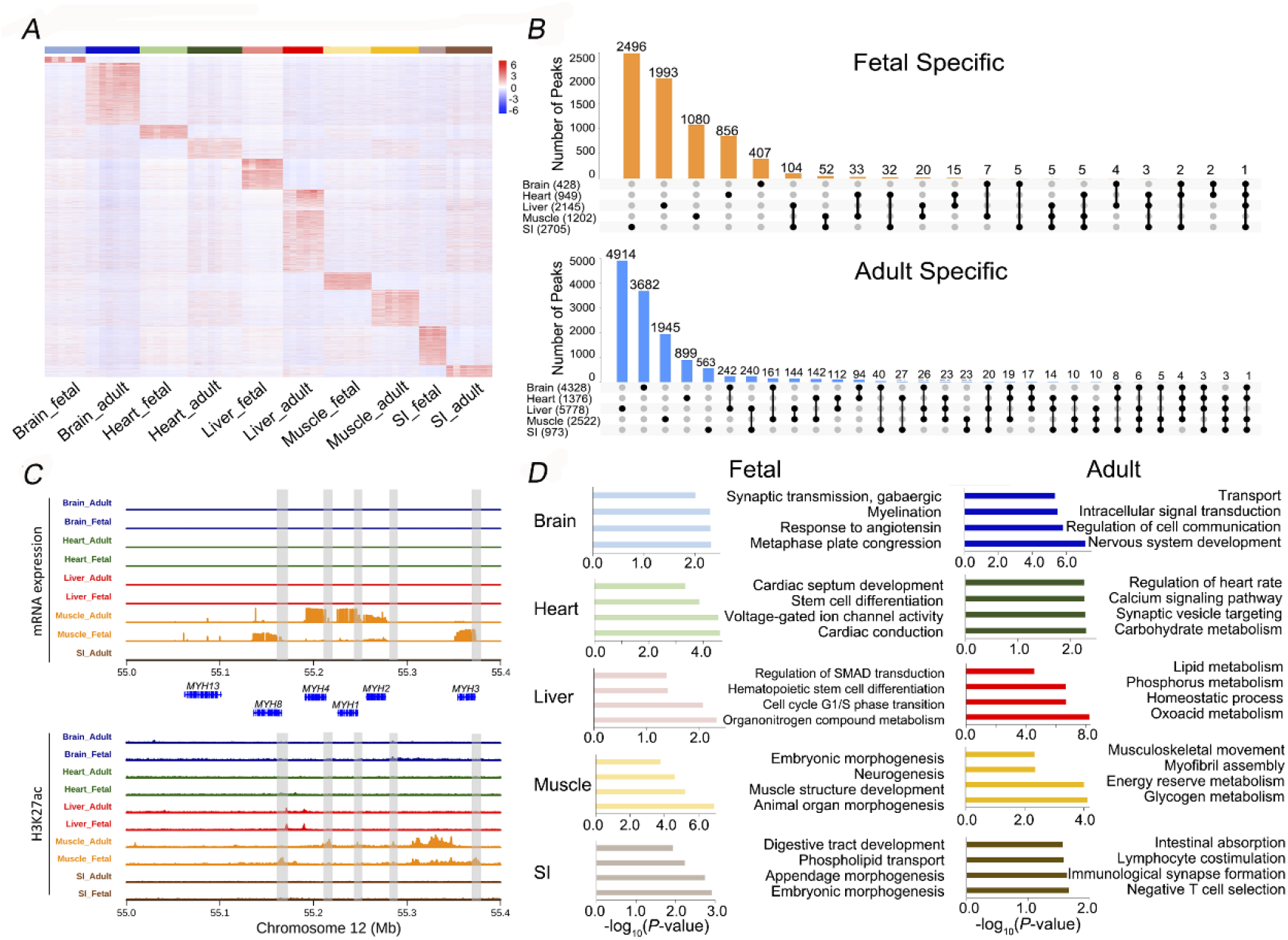
Functional annotation of developmental stage specific H3K27ac peaks in the five tissues. (**A)** Heatmap of all developmental stage specific peaks in the five tissues. Each row of the heatmap shows normalized density of read-depth at one peak. (**B)** Intervene plot showing the number of adult or fetal specific peaks that were shared across the five tissues. (**C)** The H3K27ac activity and mRNA expression tracks in the genomic region on chromosome 12 containing the myosin heavy chain gene family. (**D)** Bar plot showing representative biological processes enriched in target genes of the developmental stage specific peaks in the five tissues.

We focused on a region containing six myosin family genes (*MYH13*, *MYH8, MYH4, MYH1, MYH2* and *MYH3*) that are regulated in a developmental stage specific manner in muscle. This region was reported to be significantly associated with intramuscular fat content (IMF) in Korean native pigs, and a 6-bp deletion upstream of the *MYH3* gene was proposed to be the causative mutation (Cho et al., 2019). We observed fetal specific expression of *MYH8* and *MYH3* in muscle coinciding with fetal specific H3K27ac peaks at the promoter regions of these two genes. Stronger expression of the *MYH4*, *MYH1* and *MYH2* genes in adult muscle were associated with adult-specific peaks close to these genes (**Fig. 5C**). The fetal specific expression of *MYH3* possibly argues against *MYH3* as causal gene for the loci for IMF, or the effect of *MYH3* on the IMF could result from a mechanism operating in the fetus.

Gene ontology enrichment analysis showed that the target genes of the fetal specific peaks were enriched in biological processes like cell cycle, tissue development and other pathways that match the biology of fetal status of the corresponding tissues. For example, gene sets related to myelination (Jakovcevski et al., 2009) (*q* = 0.005) and GABAergic synaptic transmission(Kirmse et al., 2018) (*q* = 0.0098) (that are important for fetal brain development) were enriched in target genes of fetal brain specific peaks, while gene sets related to hematopoietic stem cell differentiation (*q* = 0.052) (that match hematopoiesis function of fetal liver (Popescu et al., 2019)) were enriched in target genes of fetal liver specific peaks (**Fig. 5D and Supplemental Table S8**). For adult specific peaks, we identified biological processes including choline transport (*q* = 0.007) and cardiac muscle contraction (*q* = 0.008) for heart, oxo-acid metabolic process (*q* = 5.8×10^-9^) for liver, glycogen catabolic process (*q* = 3.8×10^-4^) and skeletal muscle contraction (*q* = 0.011) for muscle, and T cell selection (*q* = 0.022) and intestinal absorption (*q* = 0.027) for small intestine (**Fig. 5D**). These results underscore the important roles of H3K27ac for the fetal to adult functional switch of the corresponding tissues through regulating the genes and pathways with developmental stage specific roles.

We also found significant overrepresentation of 140 TFBS in the developmental stage specific peaks (**Supplementary Table 9**). Many of the enriched transcription factors have physiological functions that are relevant for the biology of the cognate tissues at the respective developmental stage. The most enriched TF included FOXD3 (neural crest development), SOX17 (cell cycle in mouse brain) and SOX2 (adult mouse brain neurodegeneration) for adult brain specific peaks, and RFX3 (early brain development) for fetal brain specific peaks; GATA4 (myocardial regeneration in neonatal mice), GATA6 (cardiac development) and TBX20 (cardiac progenitor formation) for fetal heart specific peaks; HNF4a (cell differentiation and proliferation in liver), FOXK1 (liver cancer), FOXO3 (regulate hepatic triglyceride metabolism) for adult liver specific peaks, and GATA2 and GATA1 (hematopoietic factors in fetal liver cells) for fetal liver specific peaks; MEF2D, MEF2C, MEF2b, SIX1 and SIX2 that are essential for muscle differentiation and development for adult muscle specific peaks, and myogenic regulatory factors (MYF5, MYOG and MYOD) that are relevant to embryonic development for fetal muscle specific peaks; IRF2 (regulate IL-7 in intestinal epithelial cells) for adult intestine specific peaks, and HNF4a (maturation of fetal intestine) for fetal intestine specific peaks (**Supplementary Fig. 14**). Further independent analyses or experiments are required to validate these finding.

### Identifying H3K27ac peaks with differential activity between Bama Xiang and Large White pigs

To reveal peaks differing in activity between the two investigated breeds, we performed a linear regression analysis and revealed 264 peaks with significant differential activity between Large White and Bama Xiang pigs at *q* value threshold of 0.05 **(Supplementary Table 10)**. These peaks were associated with genes involved in IL-6 signaling pathway (Benjamini-Hochberg corrected *q* = 0.011), ovarian follicle development (*q* = 0.012) and response to thyroid hormone (*q* = 0.013) **(Supplementary Table 11)**. They were enriched for the binding motifs of three transcription factors including ERG (*q* = 0.016), GAPBA (*q* = 0.033) and CEBP (*q* = 0.033) **(Supplementary Table 12)**. Among the 264 peaks, 52 and 13 were consistently having higher activity in samples across all tissue-development stages in Bama Xiang and Large White pigs, respectively (**Supplementary Fig. 15A**).Two representative examples are showed in **Supplementary Fig. 15D-E**. The relevance of these peaks and their target genes with the phenotypic differences between the two breeds require further investigations.

### H3K27ac and DNA methylation at the *XIST* gene promoter region regulate X chromosome inactivation

We next investigated the impact of sex on peak activity. At q value threshold of 0.05, we identified 13 sex biased peaks, including two located on the X chromosome and 11 located on the Y chromosome (**Fig. 6A**). The H3K27ac peak region 59,307,569-59,310,238 on chromosome X coincided with promoter region of *XIST* and exhibited female specific activity across all five tissues and two developmental stages (**Fig. 6B-D**). This coincided with the female specific expression of the *XIST* gene, which is a well-studied non-coding gene driving X chromosome inactivation (Marahrens et al., 1998). As DNA methylation is highly related to H3K27ac (Charlet et al., 2016), we further examined the distribution of muscle DNA methylation based on the whole genome bisulfite sequencing in the *XIST* gene region in two female and two male adult Large White pigs(Min Zheng, 2020), and found that the peak region (chrX: 59,307,569-59,310,238) was completely methylated in males and hemimethylated in females (**Fig. 6E**). We assume that the complete methylation in males is accompanied by de-acetylation of H3K27 in the promoter region of *XIST* and silencing of the *XIST* gene, while in females, the promoter region of *XIST* from the active X chromosome is fully methylated, and the corresponding region on the inactivate X chromosome is de-methylated and associated with H3K27 acetylation and transcription of the *XIST* gene, which in turn initiates the downstream pathways of X chromosome inactivation. Taken together, these analyses support the involvement of H3K27ac and DNA methylation at the promoter region of the *XIST* gene in the regulation of X chromosome inactivation. As far as we know, this has not yet been reported elsewhere despite the fact that initiation, spreading and maintenance of X inactivation in mammals is well studied (Marks et al., 2009). Thereby the analysis herein provides complementary information about this fundamental biological process conserved across placental mammals.

**Fig. 6:**
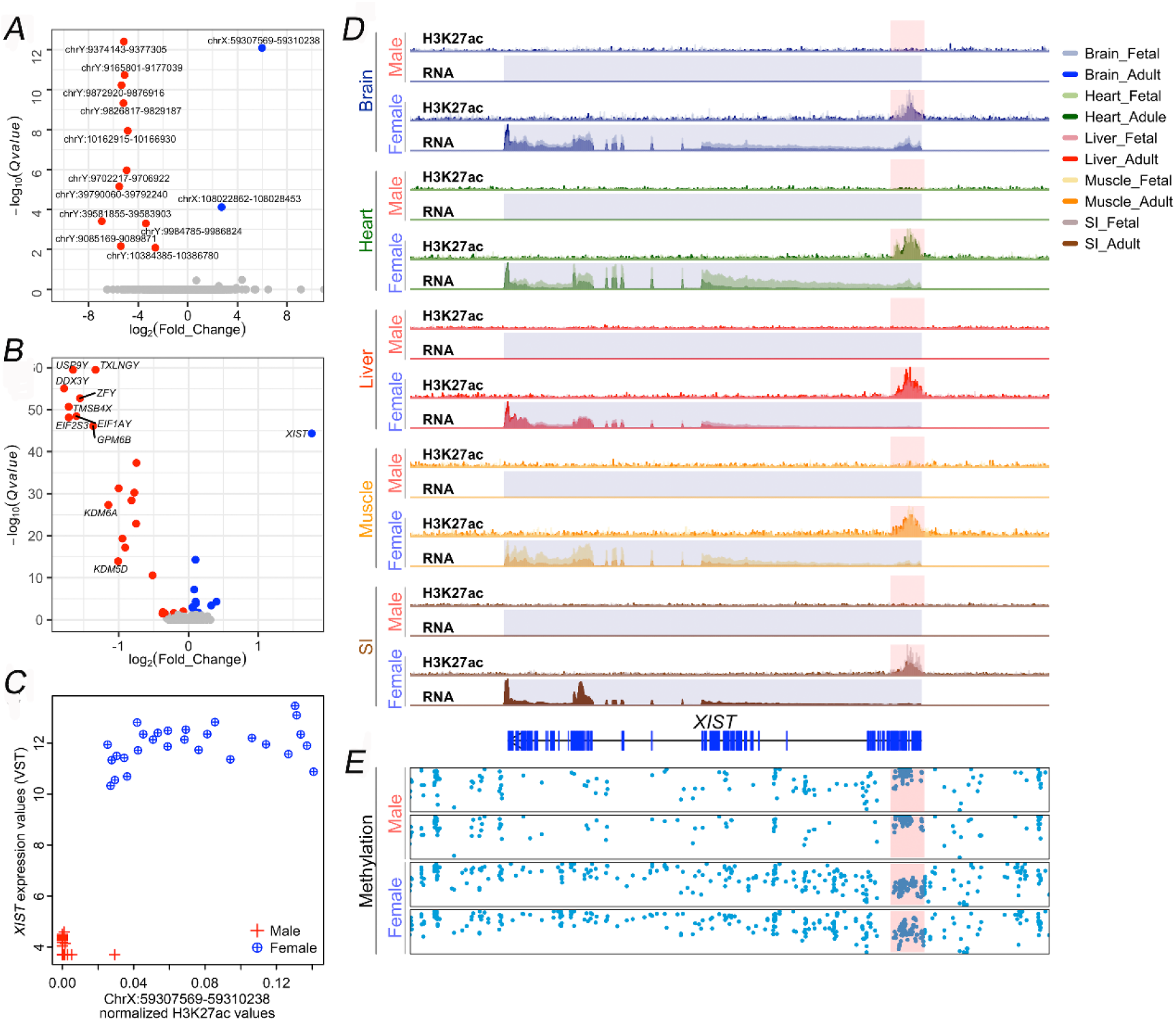
Identification of H3K27ac peak activity involved in X chromosome inactivation. (**A)** Volcano plot of peaks exhibiting significant gender specific activity. (**B)** Volcano plot of differentially expressed genes between the two genders, the annotated genes shown higher expression in males are all located on Y chromosome. (**C)** Scatter plot showing the correlation between the peak at chrX: 59,307,569-59,310,238 with expression of *XIST* gene; (**D)** Comparison of the tracks of peak activity and gene expression at *XIST* gene regions in males and females across the tissue – developmental stages; Y axis corresponding to the normalized read depths of ChIP-Seq or RNA-Seq data. (**E)** DNA methylation signals at XIST gene region in two female and two male Large White pigs. Y axis indicate percentage of reads covering corresponding sequence bases that were methylated, a dot represents a CpG island.

### Applying the H3K27ac profile to the genetic dissection of complex traits in pigs

Characterizing the regulatory elements that control relevant genes for complex traits is a key step in understanding the regulatory mechanism underlying variations in quantitative traits. Here, we investigated tracks of mRNA expression of three previously reported complex trait related genes (*PHKG1*, *LDLR* and *MSTN*) and the nearby H3K27ac landscape across the ten tissue – developmental stages. We showed that *PHKG1*, which harbors a splicing mutation associated with pH value and water holding capacity of meat (Ma et al., 2014), displayed adult muscle restricted expression that is associated with adult muscle specific activity of an H3K27ac peak in the first intron of this gene (**Supplementary Fig. 16**). The *LDLR* gene that encodes the low density lipoprotein receptor that was reported to be associated with hypercholesterolemia in pigs (Hasler-Rapacz et al., 1998), was also found to show liver biased expression associated with two liver biased active H3K27ac peaks located in the *LDLR* gene body (**Supplementary Fig. 16**). *MSTN*, a gene well-known to be associated with muscle growth in various livestock species including pigs (Aiello et al., 2018), displayed muscle specific expression that is concordant with a muscle specific H3K27ac peak in this gene (**Supplementary Fig. 16**). These examples demonstrate that the present resources provide helpful information to understand the regulatory basis of tissue specific expression of complex trait related genes.

Identifying causative mutations underlying the variations in complex traits is a key objective of genetic studies. To test whether the H3K27ac data could help to prioritize causal variants for complex traits, we examined the genomic region harboring a well-known causal mutation (chr2:1483817) in the first intron of the *IGF2* gene that increases muscle growth in pigs (Van Laere et al., 2003). We found that the causal mutation locates within a H3K27ac peak that shows greater activity in fetal liver and muscle, correspondingly the *IGF2* gene also displayed higher expression in fetal liver and muscle (**Fig. 7A-B**). The causal mutation is known to abrogate the binding of an inhibitor that is normally expressed in postnatal skeletal muscle. It is interesting to notice that the exact position oif the mutation coincides with an ∼300 bp trough in the H3K27ac peaks in fetal muscle and liver. The precise meaning of this observation remainsd to be elucidated (**Fig. 7C**). Overall, these results suggest H3K27ac signals generated hereby are valuable resource that is helpful to prioritize causal variants and understand the regulatory mechanism of causal genes for quantitative traits in pigs.

**Fig. 7:**
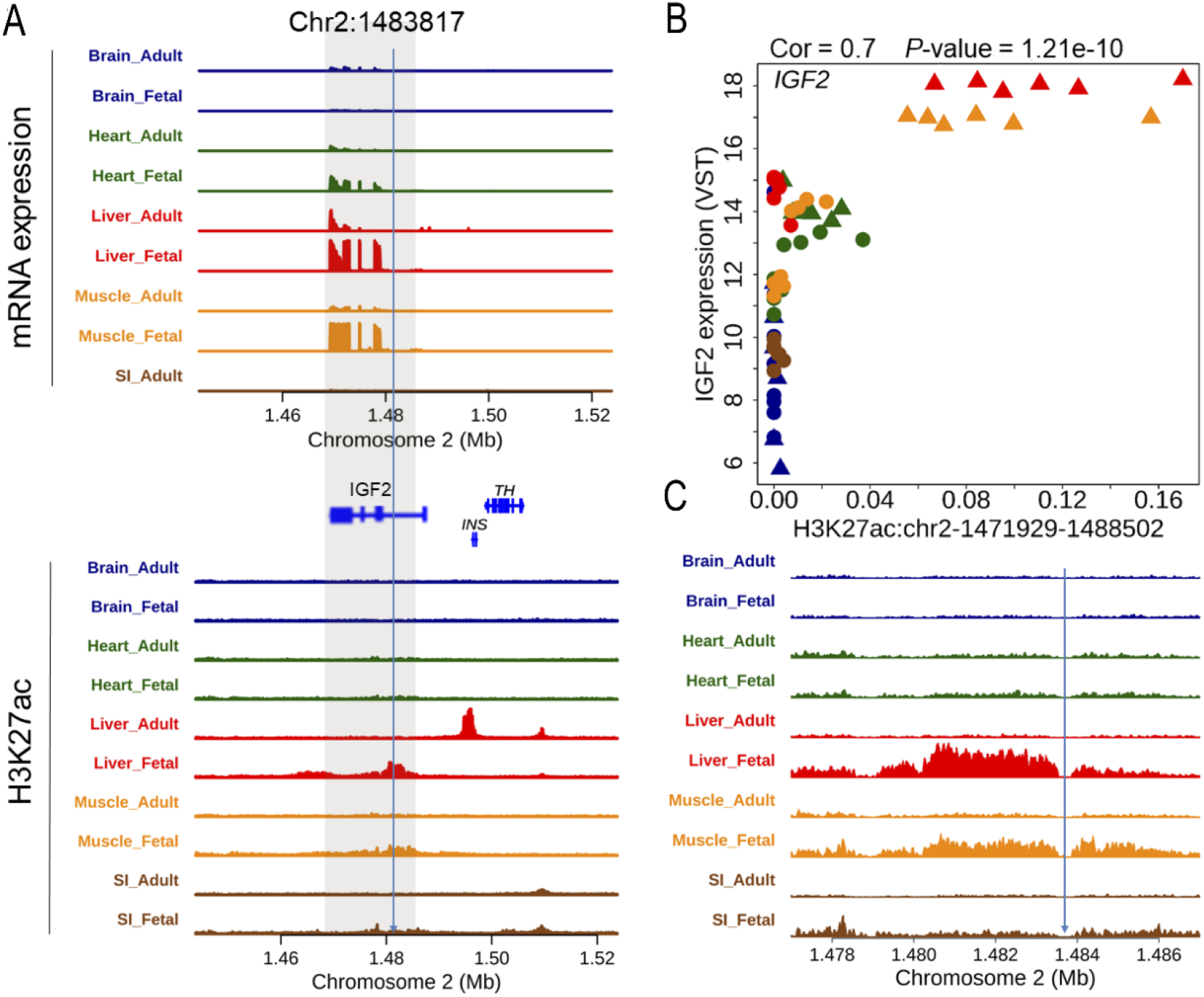
Prioritizing causative mutation using H3K27ac ChIP-Seq data. (**A)** The causal mutation for muscle growth at first intron of *IGF2* (chr2:1483817) genes located within an H3K27ac peaks in first intron of IGF2 gene exhibiting greater activity in fetal liver and muscle in pigs; **(B)** Correlation of the intronic peak harboring the causal mutation with expression of IGF2 across samples; **(C)** A zoom in plot of the H3K27ac ChIP-Seq track reveal a foot print of transcription factor binding site around the causal mutation.

### A web-based browser for easy access to the H3K27ac peak and gene expression data

To improve the accessibility of the data and results generated in this study, we developed a web-based browser that allows fast navigation of 1) the H3K27ac ChIP-Seq and RNA-Seq read coverage of the ten different tissue-developmental stages across the genome, and 2) the links between H3K27ac peaks and nearby genes based on significant correlations. The browser is largely self-explanatory and can be easily customized to visualize the subset of ChIP-Seq or RNA-Seq tracks. The colors of the tracks can be manually specified. Moreover, the visualized tracks can be saved as high resolution publishable figures. We expect that these attributes of the browser will be particularly helpful for colleagues who are interested in looking at candidate regions that associate with a certain complex trait or regions that are subject to artificial or natural selection (http://39.108.231.116/browser/?genome=susScr11).

## Discussion

Active promoters and enhancers, marked by the H3K27ac chromatin modification, are important sequences that regulate gene transcription (Creyghton et al., 2010). H3K27ac marks have a highly dynamic activity across different tissues (Roadmap Epigenomics, et al., 2015) and development stages (Gorkin et al., 2020) and are enriched in disease associated variants (Roadmap Epigenomics, et al., 2015). Until now, H3K27ac in different tissues and developmental stages has remained unexplored in the pig. In this study, we present the largest spatiotemporal spectrum of H3K27ac activity to date, in samples covering five tissues, both prenatal and postnatal stages, two breeds with divergent evolution history and phenotypes, and the two genders, hence largely expanding the database of H3K27ac regions in pigs. There are several lines of evidence that support the quality of our H3K27ac ChIP-Seq data. These include 1) the high correlation of peak activities between technical replicates (**Supplementary Fig. 17**), 2) the high proportion of liver peaks overlapping with those reported in independent studies, 3) the highly significant correlation of proximal peaks with their cognate genes within each sample, 4) the accurate classification of samples by their tissue and developmental stage based on the quantified peak activity.

A total of 62 samples covering all tissues, ages, sexes and breeds were simultaneously analyzed for both H3K27ac ChIP-Seq and RNA-Seq, allowing us to reveal the landscape of association between H3K27ac activity and gene expression. Notably, we observed considerable numbers of significant negative peak-gene correlations, which showed different distribution in terms of peak-gene distances compared to those positive peak-gene correlations, reflecting a different regulatory mechanism between positive and negative peak-gene correlations. The observed negative peak-gene correlations could also be attributed to the effects of TFs or other histone modifications that had not been arrayed. Studies have been implemented to identify silencers through high-throughput screening genomic sequences that repress the transcription of cell death protein (Pang and Snyder, 2020), or correlating the H3K27me3-DNase I hypersensitive site (DHS) peaks with nearby genes across diverse cell lines (Huang et al., 2019). Nevertheless, very few studies have reported negative correlations of H3K27ac activity with gene expression. Further experimental validation is required to confirm the assumption of silencing effect of H3K27ac on gene transcription.

We revealed a small proportion (2.6%, 2679 in 101,290) of peaks that show constitutive activities across the ten tissue-developmental stages, most of which are proximal peaks. These peaks were demonstrated to be associated with the high and stable expression of genes across tissue - stages, making the integenic or intronic regions close to ubiquitously active H3K27ac good candidates safe harbor regions for genomic engineering in pigs. Currently, studies to characterize the safe harbors in pigs focused on well-known house keeping genes such as GAPDH (Han et al., 2019) and ACTB (Xiong et al., 2020). This study provides more than hundreds candidate safe harbors containing genes that showed higher and more stable expression compared to two of the known safe harbors (H11 and Rosa26), hence largely expanding the candidates safe harbor regions in pigs.

Consistent with the reports in humans and mice that the enhancers marked by H3K27ac show highly variable activity across tissues and developmental stages (Gorkin, et al., 2020, Nord, et al., 2013, Roadmap Epigenomics, et al., 2015), we also identified a large number of tissue and developmental stage specific distal peaks. We found that these peaks were enriched for binding sites of transcription factors and regulate the genes with function matching the biology of corresponding tissue-stage, such as transcription factors and gene pathways related to hematopoiesis in fetal liver and immune response in adult small intestine, supporting that the ChIP-Seq and RNA-Seq data covering different tissues and developmental stages provide a valuable resource that is helpful to understand the regulatory mechanism underlying the tissue differentiation and development of pigs. The developed browsers further improve the accessability of the data and results generated in this study.

The analysis of sexually differential activity suggested that involvement of H3K27ac and methylation at promoter region of *XIST* gene in regulating the expression of *XIST*, which in turn initiates the pathways underlying X chromosome inactivation. A number of studies were performed to characterize the downstream processes regulated by XIST genes, such as histone modifications along the X chromosome (Zylicz et al., 2019) and gene expression patterns such as genes escape from X chromosome inactivation (Tukiainen et al., 2017). To our knowledge, this is the first study to report the association of histone acetylation and DNA methylation with XIST gene expression, and in turn X chromosome inactivation in pigs, and very few if any studies have reported the links between H3K27ac and DNA methylation with regulation of *XIST* gene in humans and other model organism.

We also explored the utility of the data to interpret the genes and mutations associated with economically important traits in pigs. We showed that the expression of trait associated genes such as *MYH3*, *PHKG1*, *MSTN* and *LDLR* were correlated with H3K27ac peaks, and displayed tissue or stage specific expression, enhancing the knowledge on the regulatory basis of these trait associated genes. Moreover, we also showed a proof of principle example that the well-known causal mutation for muscle growth was located in a peak showed higher activity in fetal muscle and liver, which associated with higher expression of *IGF2* gene in fetal muscle and liver tissues, agreeing with a recent report on the gene expression pattern of *IGF2* gene in mice (Younis et al., 2018). These results demonstrate that the data is valuable to characterize regulatory pattern of candidate genes and prioritize the causal variants for complex traits in pigs. However, extensive intersecting of the peaks identified hereby with GWAS loci for various traits in pigs e.g., from pig QTL database (Hu et al., 2013) generated few sensible enrichments (data not shown). We reasoned that large proportions of association studies in pigs were currently based of SNP array in single or limited number of populations, many identified loci reside in regions with high linkage disequilibrium (Gong et al., 2019, Zhang et al., 2017), which present a challenge to enrichment analysis of traits associated loci. We expect that, with the growth of genome sequence based GWAS loci catalogues in pigs, our data will provide a valuable map to characterize the regulatory mutation underlying complex traits in pigs.

## Materials and methods

### Samples

Five tissues, including brain (cortex), heart (left ventricle), liver, small intestine (duodenum), skeletal muscle (longissimus dorsi) were collected from seven fetuses (two males and two females from Large White at day 75 and two males and one female for Bama Xiang at day 74 post insemination), and eight adult pigs (two males and two females from Large White at day 150 and two males and two females from Bama Xiang at day 132-141 postnatal) (**Fig. 1 and Supplementary Table 1**). We carefully dissected tissues following the standardized sample collection protocols of the FAANG Project (https://www.faang.org/bbs?s=protocols..txt). Tissues were flash frozen in liquid nitrogen immediately after collection, and then stored at temperature of −80 °C.

These tissues develop from the three germ layers: ectoderm (cortex of brain), mesoderm (left ventricle of heart and *longissimus* muscle) and endoderm (liver and duodenum), and are relevant to meat production and biomedical research. Brain is frequently studied for dissecting psychiatric and mental disorder in humans (Dedova et al., 2009), and also potentially useful in studying pig behavior. Heart serves as a pump to circulate the blood that is essential for survival, and pig is a potential heart donor for humans (McGregor and Byrne, 2017). *Longissimus* muscle mass is directly relevant to meat production. Liver is central to lipid and protein metabolism and detoxification, and plays a major function in fetal hematopoiesis (Zhao and Duncan, 2005). The small intestine is an organ for food digestion, and may affect feed conversion in pigs (Vigors et al., 2016). The inclusion of two developmental stages allows investigation of regulatory basis underlying the functional switch of a tissue from prenatal to postnatal stage. Moreover, the two breeds, Large white and Bama Xiang, represent the two major swine domestication centers (Near east and China) (Larson et al., 2005), and differ considerably for a variety of phenotypes including body size, coat color and fat deposition. The wild ancestors of the two breeds diverged about 1.2 million years ago (Frantz et al., 2013).

### H3K27ac ChIP-Seq

The chromatin immunoprecipitation of samples was processed using the SimpleChIP Plus Enzymatic Chromatin IP Kit (Magnetic Beads) (CST Danvers, MA, 01923, United States). Briefly, about 200-300 mg sample was minced in 1 ml PBS, cross-linked using 37% formaldehyde at room temperature for 10 minutes. The cross-linked sample was terminated by 10x glycine for 5 minutes and lysed with buffer A and B in the IP kit. The DNA was then sheared to target size of 100-300 bp by sonication in 500 μl ChIP buffer in the IP kit, 10 μl of the DNA solution was kept as input. ChIP was performed using 5 μg H3K27ac antibody (Active Motif Carlsbad, CA 92008, North America) overnight. The DNA bound by the antibody were enriched using ChIP-Grade Protein G Magnetic Beads in the IP kit, and then incubated with 2ul 20mg/ml proteinase K and 6 μl 5 M NaCl with at 65 °C to reverse cross-links. The immunopreciptated DNA was then extracted. DNA sequencing was performed on Illumina HiSeq 2500 in a single read 50bp run on Illumina HiSeq 2500, the raw sequence reads were filtered by removing whole reads with 1) contaminated adaptor sequence; 2) with more than half of bases with Phred quality score < 19; 3) with > 5% of ambiguous/undetermined (N) bases.

### H3K27ac peak calling and processing

The clean reads were mapped to the pig reference genome Sscrofa 11.1 using Burrows-Wheeler Aligner (BWA) (Abuin et al., 2015), allowing two mismatches. MACS (Model-based Analysis for ChIP-Seq) version 2.1.0 peak caller was used to infer peaks in each individual with option of “--broad and –broad-cutoff 0.1” (Zhang et al., 2008). Bedtools was used to combine the peaks identified in each individual to a reference peak set, by ignoring the 5% largest peaks with peak length larger than 13.1 kb, as we considered that these large peaks could be regions of multiple adjacent peaks, for which we could not quantify their individual activity accurately. The extent of developmental stage-specific peaks that were shared across tissues were visualized using Intervene program (Khan and Mathelier, 2017).

### Funtional annotations of ChIP-Seq peaks and Gene ontology enrichment analysis

The pig whole genome GERP score was downloaded from Ensembl Database, we computed the GERP score of a peak by averaging the GERP scores of bases within that peak. GC-content of a peak was caculated using EMBOSS geece software (Rice et al., 2000). The enrichment of different genomic features and TFBS were conducted using HOMER program. The gene ontology enrichment analysis was performed using ClueGO (Bindea et al., 2009). The *P* values for the enrichment of GO terms were corrected using Benjimini-Hochberg approach.

### RNA-Seq

Total RNA was isolated from tissues using Trizol reagent (Invitrogen, USA), mRNA was enriched using magnetic beads attached with poly-T oligo. The cDNA were then synthesized using random hemamer primers. The cDNA was then purified, ligated to index adapters, size selected using AMPure XP beads, and then PCR amplified to generate the strand specific cDNA libraries. The library was sequenced through paired end 150 (PE 150) run on Illumina HiSeq 4000. The raws reads comtaminated with adapter sequences and with 1) more than 50% of low quality bases (Q < 19) and 2) percentage of N bases > 10% were removed. The clean reads were mapped to pig reference genome Sscrofa 11.1 using STAR-2.5.3a (Dobin et al., 2013). The transcripts were assembled and merged with Stringtie by referring to Ensembl GTF (98.111), which were further merged with Ensembl GTF (98.111) to obtain a customized GTF file as the reference for gene expression quantification, which contains a total of 44415 genes/transcripts that consist of 24129 protein coding genes, 10060 de novo assembled genes or transcripts, 6985 long noncoding RNA, 1593 pseudogenes, 474 snoRNA, 440 miRNA and 773 other transcripts. The expression levels of genes were quantified using FeatureCounts 1.5.3 (Liao et al., 2014). We kept genes supported by at least 500 counts in 62 RNA-Seq samples. After removing the mitochondrial genes, variance stabilizing transformation (VST) algorithm from DESeq2 (Love, et al., 2014) was used to normalize the expression values for each sample.

### Statistical analyses

Unless otherwise stated, most of the statistical analyses were performed in R program version 3.5.1. The contribution of tissue, development stage, breed and sex to the variance of quantitative peak activities and gene expressions were quantified by a linear mixed model(Hoffman and Schadt, 2016). In the correlation analyses of H3K27ac activity with gene expression traits, the H3K27ac peak activity and gene expression traits were firstly adjusted for breeds, developmental stages, sex using lm function in R, the spearman correlations between the residuals of peak activity and nearby genes were computed using R program. We consider most of these long distance peak-gene correlation could be background noise, despite that part of them could reflect long distance enhancer-promoter interactions. To obtain a conservative estimate of peak-gene correlations, for each peak, we pruned the number of significant correlations identified within 500 kb by the number of significant peak-gene correlations identified within 1.5 – 2 Mb regions from corresponding peak, and finally obtained 41,105 significant correlations (35,644 positive and 5,461 negative). To identify peaks showing differential activity between two breeds, we used lm function in R by adding tissue, developmental stage and sex as fixed effects. Similar analysis was performed to assess the effect of sex on peak activity. The q values corresponding to the P values from the correlation and lm analyses were determined using R package qvalue (Storey JD, 2020).

### Population genetic analysis

The clustering of the samples based on peak activity and gene expression were performed using hcluster function with ward.D method in R program, and visualized using R package GGraph.

## Compliance and ethics

We declare that there is no conflict of interest associated with this publication.

## Acknowledgements

We are grateful to colleagues in State Key Laboratory of Pig Genetic Improvement and Production Technology, Jiangxi Agricultural University for sample collection. This work was supported by the National Natural Science Foundation of China (31790413 and 31760657).

## Author contributions

LSH designed this study, wrote and edited the manuscript; BY analysed the data and wrote the manuscript; MG wrote part of manuscript and commented on the results; YLZ and ZMZ performed the experiments, analysed the data and wrote the manuscript; TH analyzed the data; ZZ performed the experiment; WBL, ZQL, TJ, SYY, JWY and YYX assisted with experiments and data analysis.

