## Supplemental Data 1 for "The swine spatiotemporal H3K27ac spectrum provides novel resources for exploring gene regulation related to complex traits and fundamental biological process"

The PDF file includes:

Supplementary figures: Fig. S1 – S17

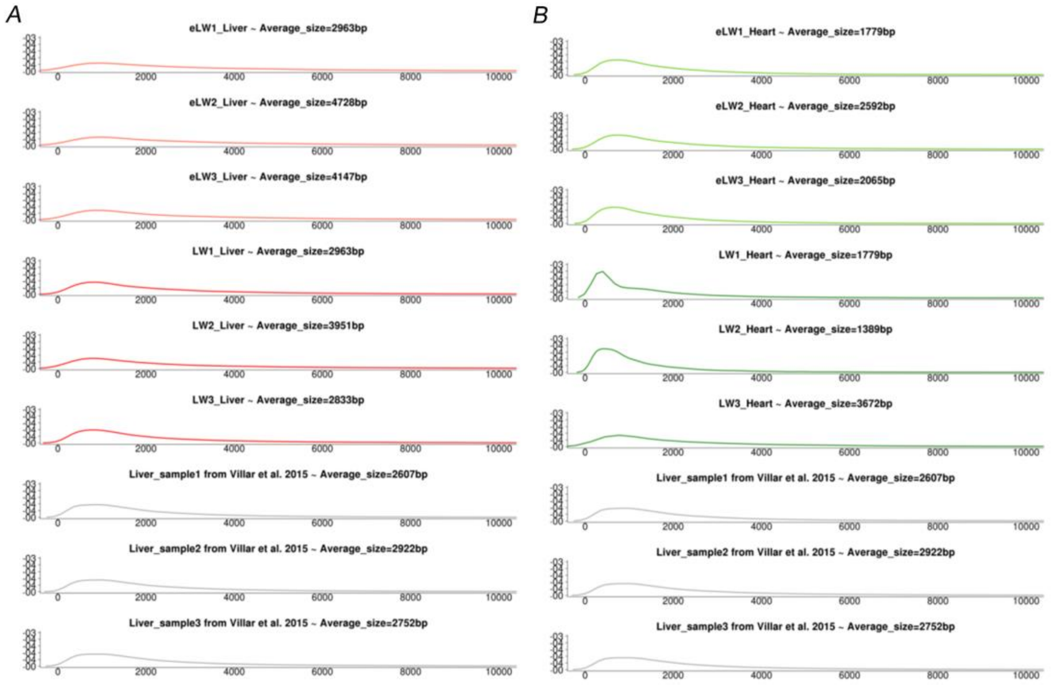


Supplementary Fig. 1: Comparison of peak size distribution and average obtained in this and previous studies[7].

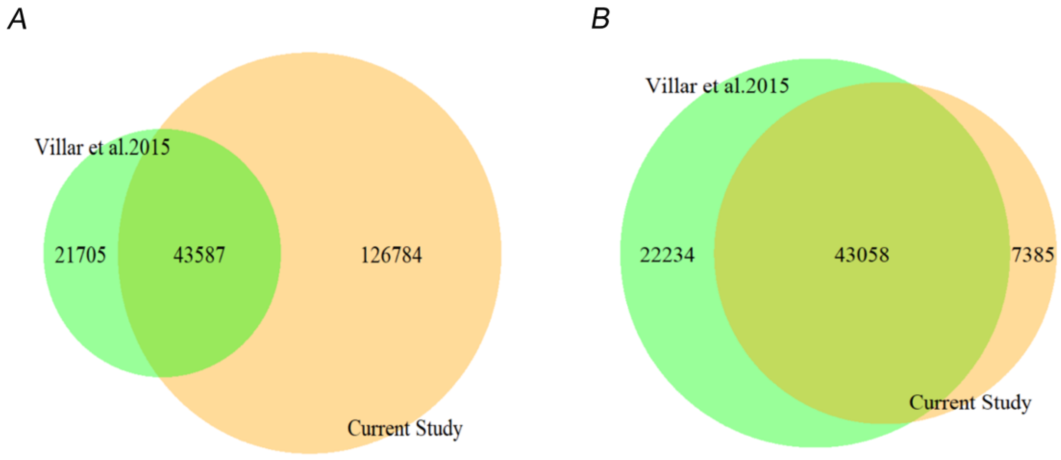


Supplemental Fig. 2: Overlap of peaks identified in this study with those previously reported[7]. (A) Venn plot showing the overlap of 170,371 peaks identified in this study with those reported in Villar et al. 2015 (B) A total of 43058 (85.4%) peaks active in at least one adult liver sample of male individual in this study overlapped with liver peaks identified in Villar et al. 2015.

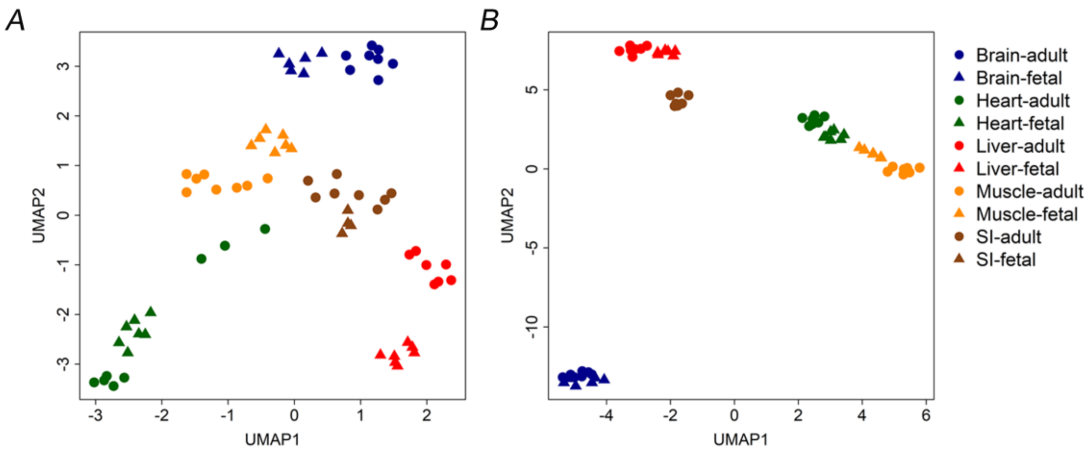


Supplemental Fig. 3: Uniform manifold approximation and projection (UMAP) plot of samples based on quantitative activity of 170,371 peaks (A) and 20319 genes/transcripts(B).

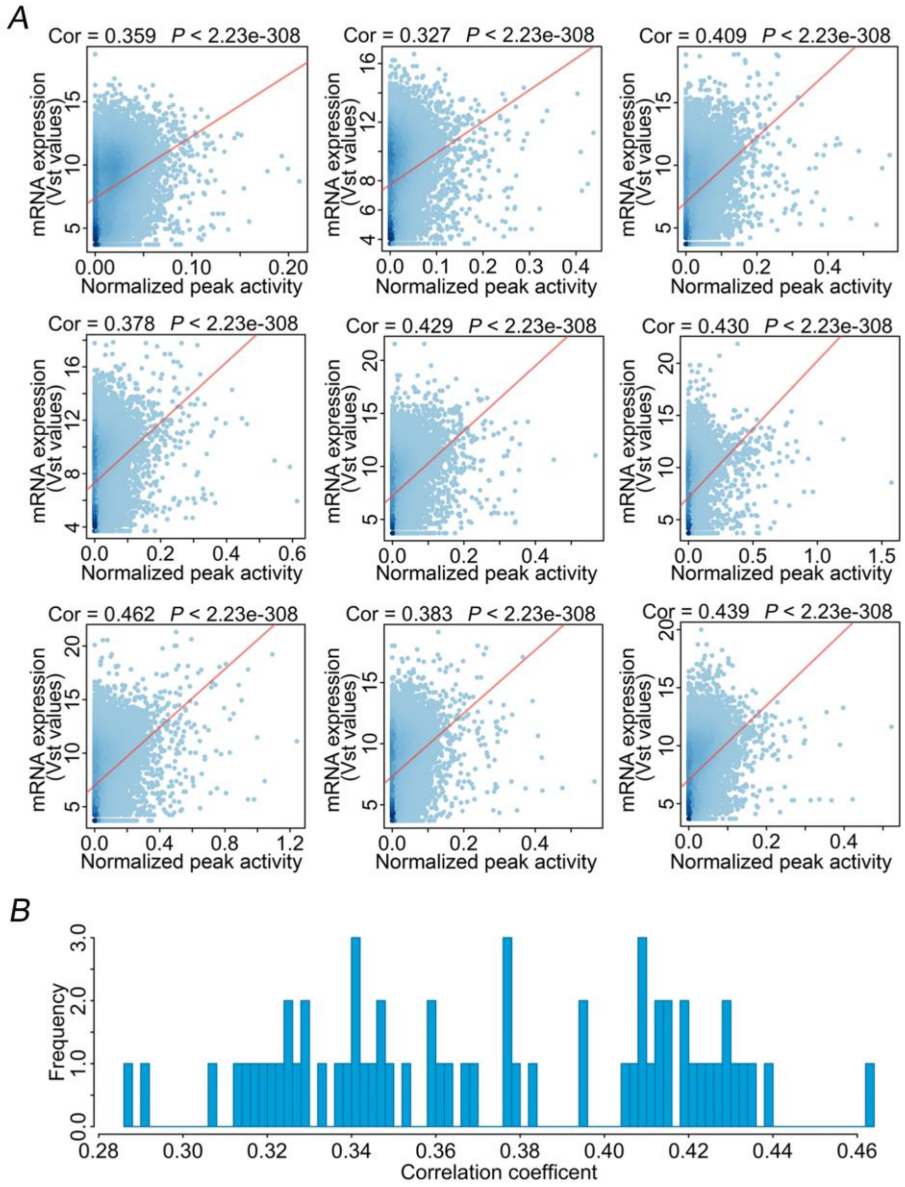


Supplemental Fig. 4: Correlation of genes and activity (measured by average depths per million reads) of their proximal peaks within each sample. (A) Scatter plots proximal peak activity and their downstream genes in 9 representative samples (average Pearson correlation coefficient r = 0.37, *P* value < $1\times10^{-300}$). (B) The distributions of correlation coefficient between peak activity and gene expression level.

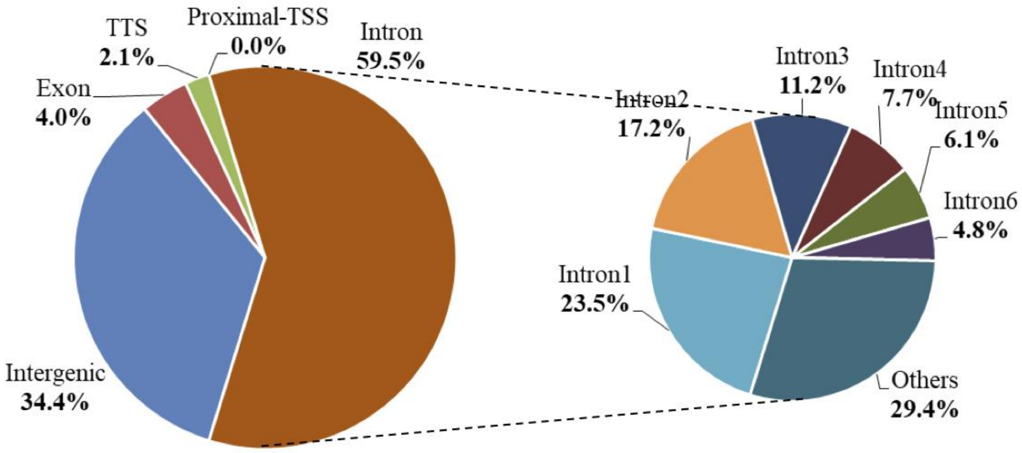


Supplemental Fig. 5: Annotating genomic features of the 82,769 distal peaks. The pie chart on the right represents the distribution of peaks located in introns, TTS, transcription termination site.

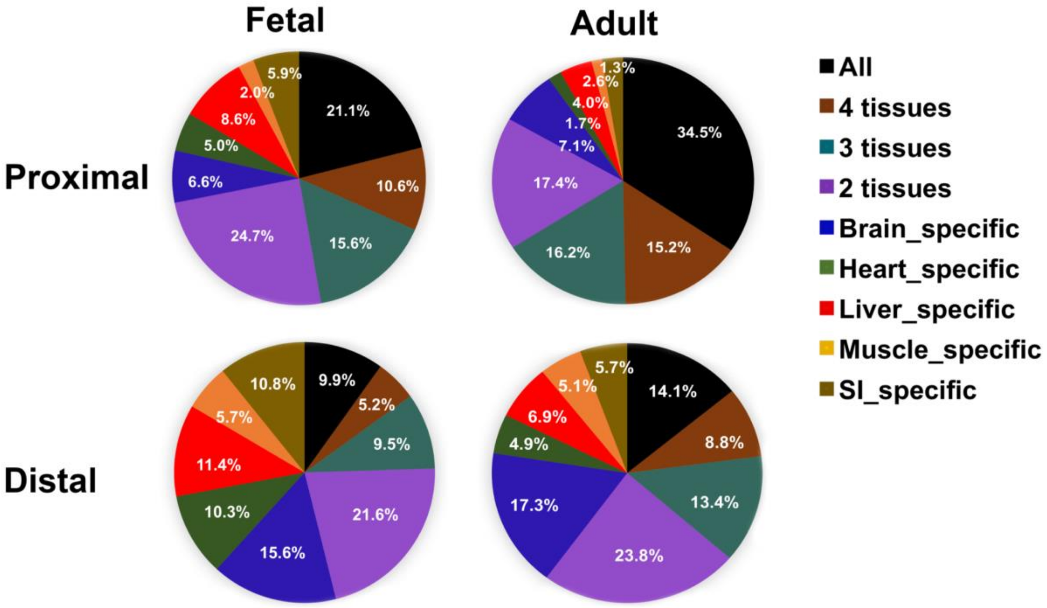


Supplemental Fig. 6: Pie plots showing the tissue specificity for proximal and distal H3K27ac peaks for fetal and adult samples.

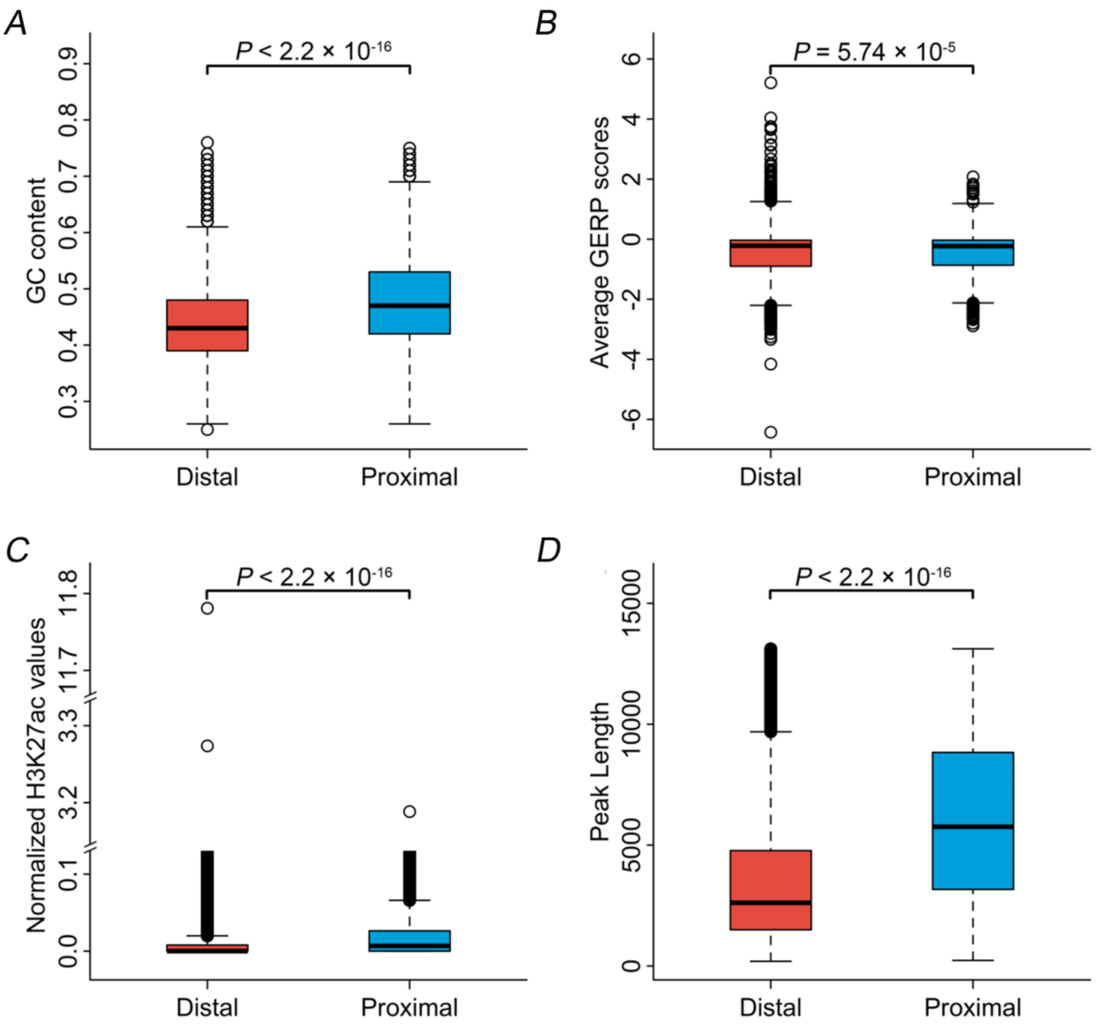


Supplemental Fig. 7: The comparison between distal and proximal peaks in terms of GC content, sequence conservation across species (GERP score), peak size and peak activity.

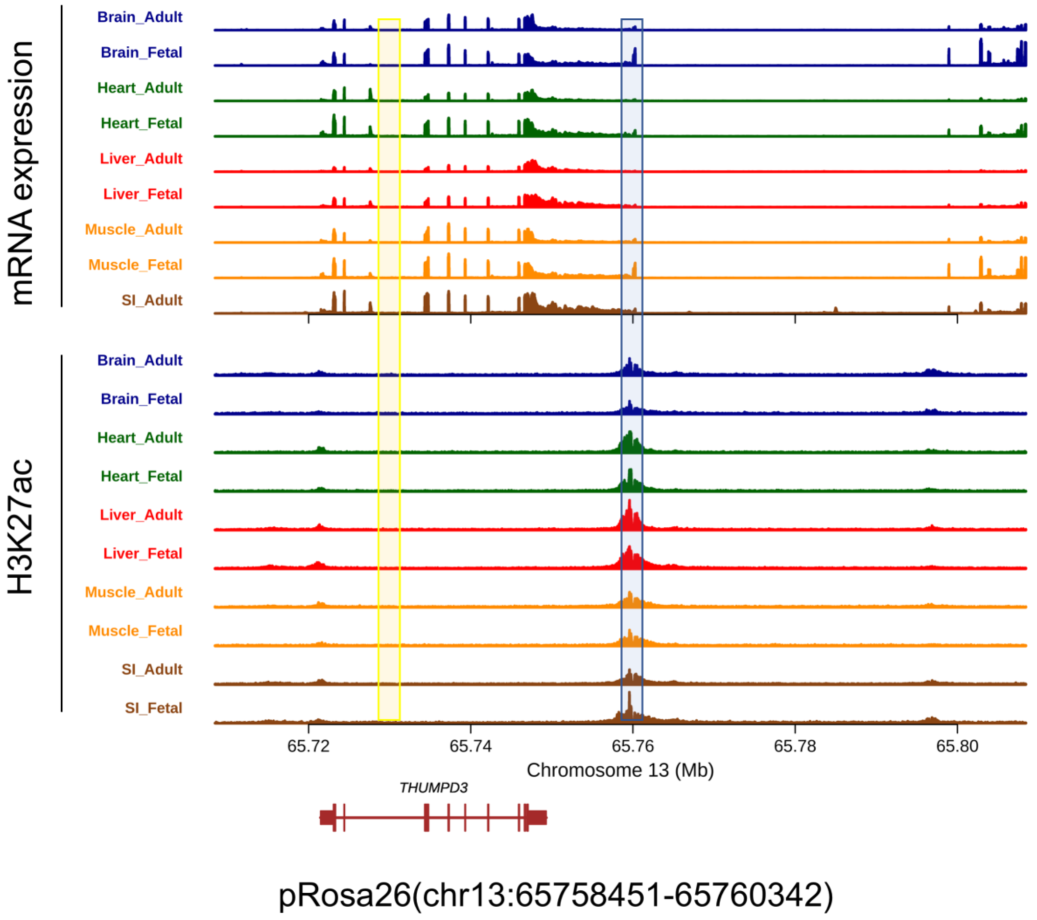


Supplemental Fig. 8: The tracks of gene expression and H3K27ac activity at Rosa26 safe harbor locus on pig genome across the tissue – developmental stages. The highlighted areas in blue represent sites of H3K27ac modification and the highlighted areas in yellow represent potential safe harbor locus.

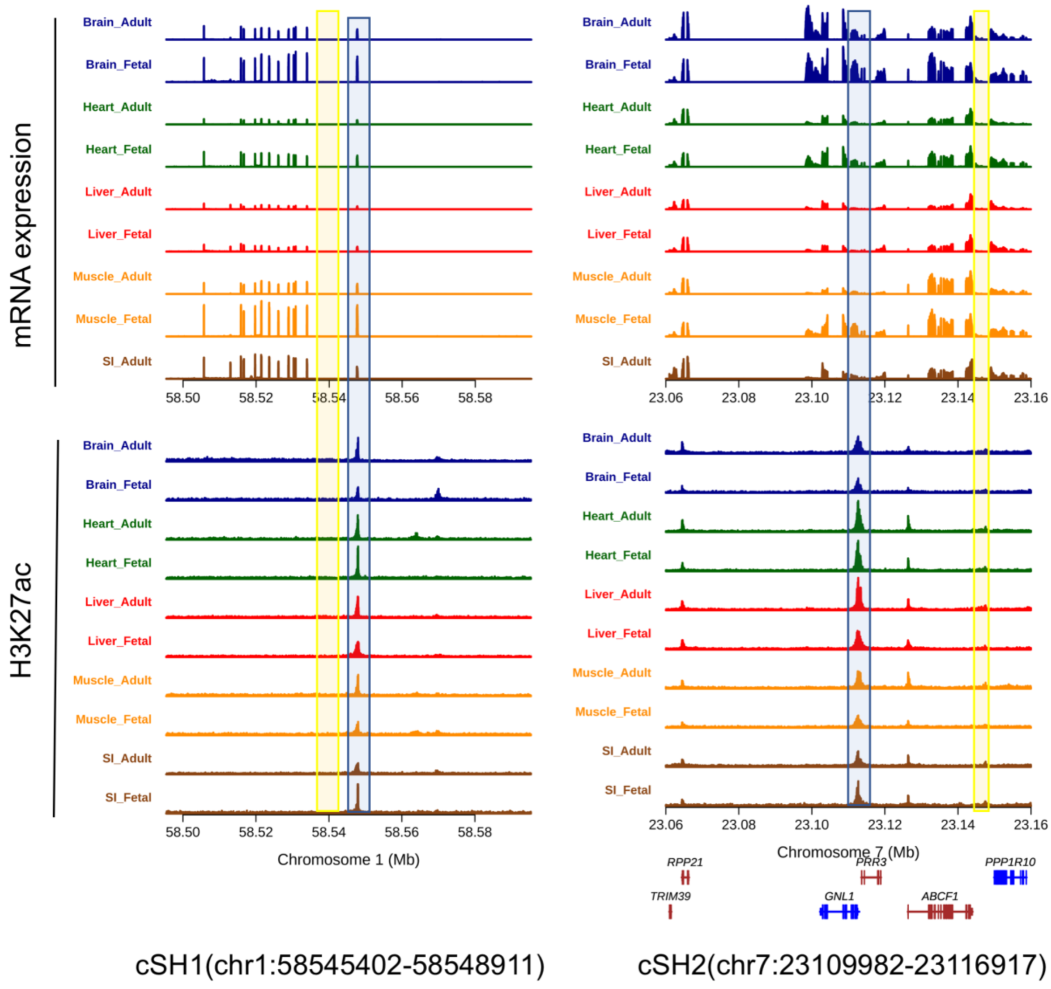


Supplemental Fig. 9: The tracks of gene expression and H3K27ac activity at two newly identified candidate safe harbor regions across the tissue – developmental stages. The highlighted areas in blue represent sites of H3K27ac modification and the highlighted areas in yellow represent potential safe harbor locus.

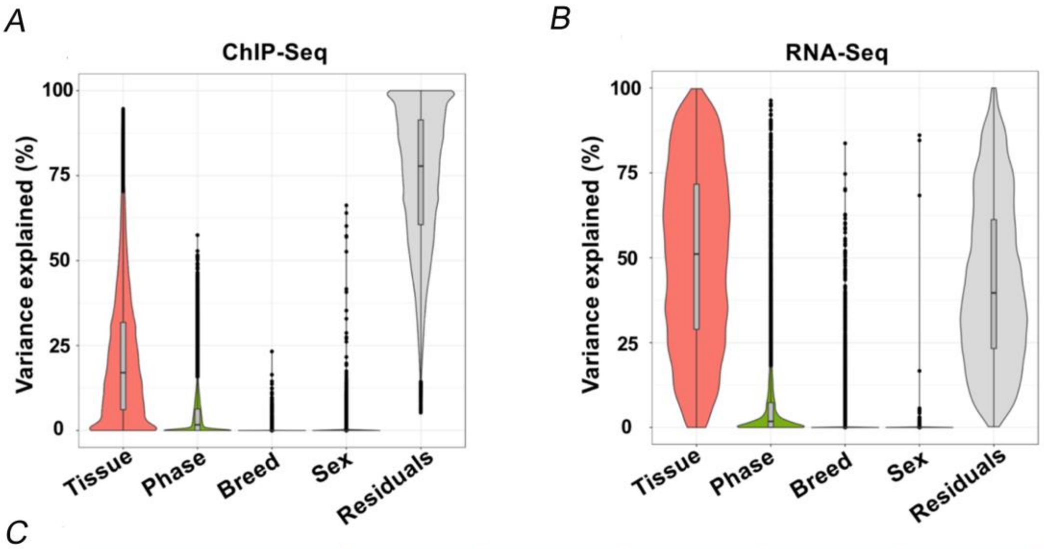


C

| Variance explained | | Tissue | Phase | Breed | Sex | Residuals |
|---|---|---|---|---|---|---|
| ChIP-Seq | Median | 0.1702 | 0.0169 | 0.0000 | 0.0000 | 0.7780 |
| | Mean | 0.2125 | 0.0398 | 0.0001 | 0.0035 | 0.7442 |
| RNA-Seq | Median | 0.5109 | 0.0179 | 0.0000 | 0.0000 | 0.3965 |
| | Mean | 0.5025 | 0.0613 | 0.0094 | 0.0002 | 0.4267 |

Supplemental Fig. 10: The contribution of tissue, developmental stage, breed and sex on the variance ChIP-Seq peak activity and gene expression traits estimated by a linear mixed model. (A) Violin plot showing distribution of proportion of variance in the ChIP-Seq peak activity explained by the tissue, developmental phase, breed and sex (B) Violin plot showing distribution of proportion of variance in the gene expression traits explained by the tissue, developmental phase, breed and sex. (C) The median and mean of the variance explained by tissue, phase, breed and sex for ChIP-Seq peaks and gene expression traits.

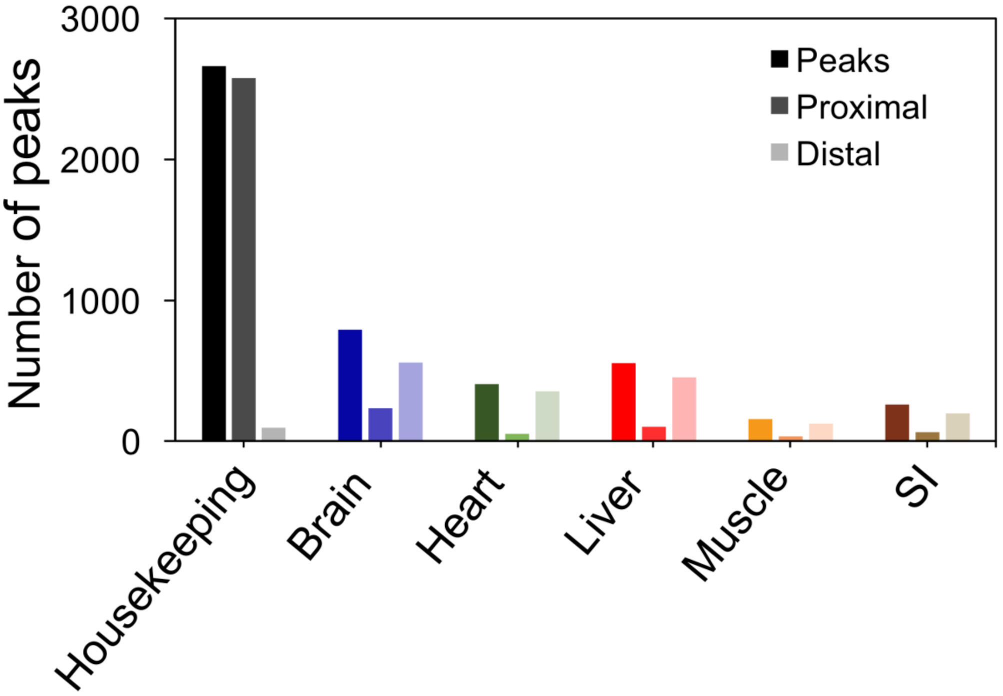


Supplemental Fig. 11: Distribution of number proximal and distal peaks in the ubiquitously active and tissue specific H3K27ac peaks.

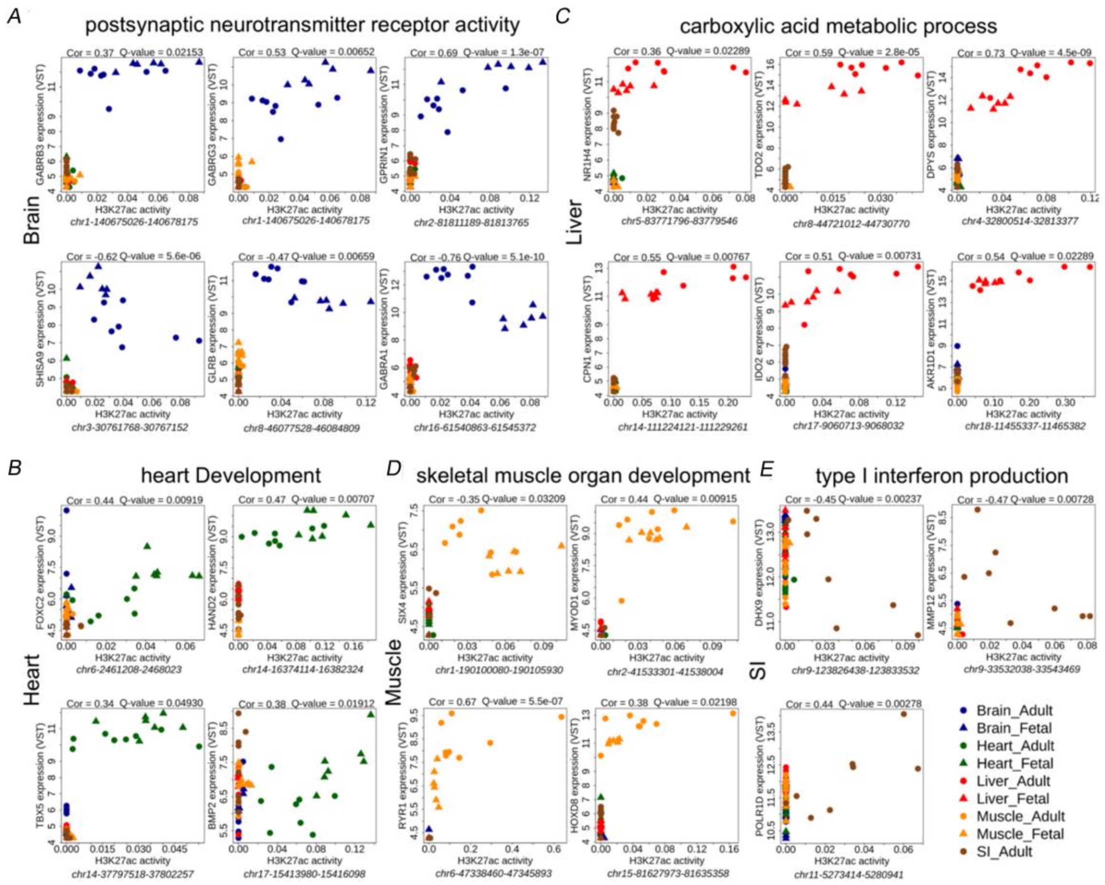


Supplementary Fig. 12: Correlation of tissue specific peaks with genes that involved in tissue specific biological pathways in the five tissues.

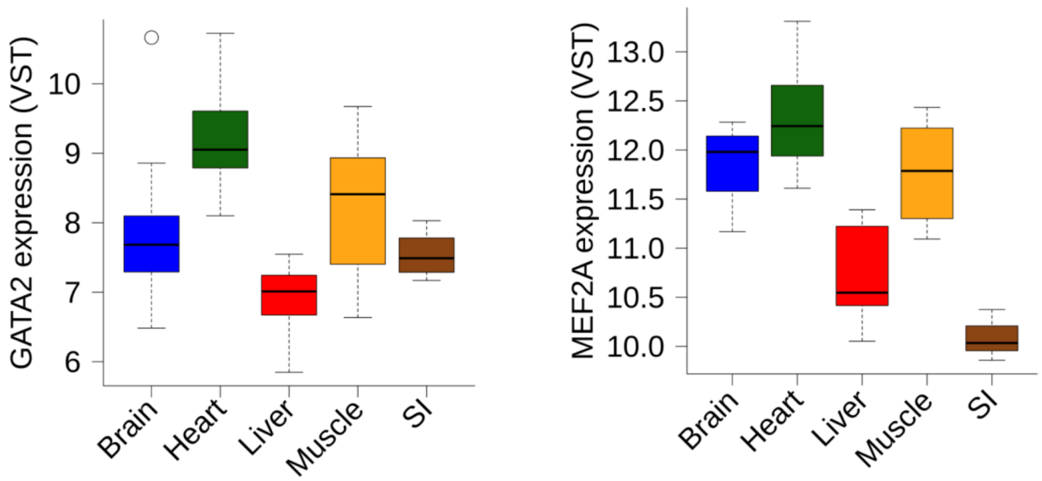


Supplementary Fig. 13: The distributions of expression of two transcription factors (GATA2 and MEF2A) with their binding motif enriched in heart

A

| Group | Motif | Name | q-value (Benjamini) | % of Peaks with Motif |
|---|---|---|---|---|
| Brain_Fetal | 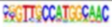 | RFX3 | 0 | 3.82% |
| Heart_Fetal | 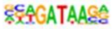 | GATA4 | 0.0001 | 18.92% |
| Heart_Fetal | 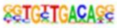 | TBX20 | 0.0001 | 4.73% |
| Heart_Fetal | 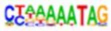 | MEF2A | 0.0172 | 8.45% |
| Liver_Adult | 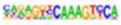 | HNF4A | 0 | 6.63% |
| Liver_Adult | 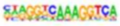 | PPARA | 0.0001 | 14.33% |
| Liver_Adult | 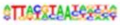 | NFIL3 | 0.0025 | 9.61% |
| Liver_Fetal | 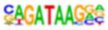 | GATA1 | 0 | 9.73% |
| Muscle_Adult | 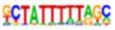 | MEF2D | 0 | 5.95% |
| Muscle_Fetal | 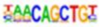 | MYF5 | 0 | 16.12% |
| Muscle_Fetal | 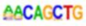 | MYOG | 0 | 19.69% |
| Muscle_Fetal | 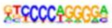 | EBF1 | 0.0474 | 12.98% |

B

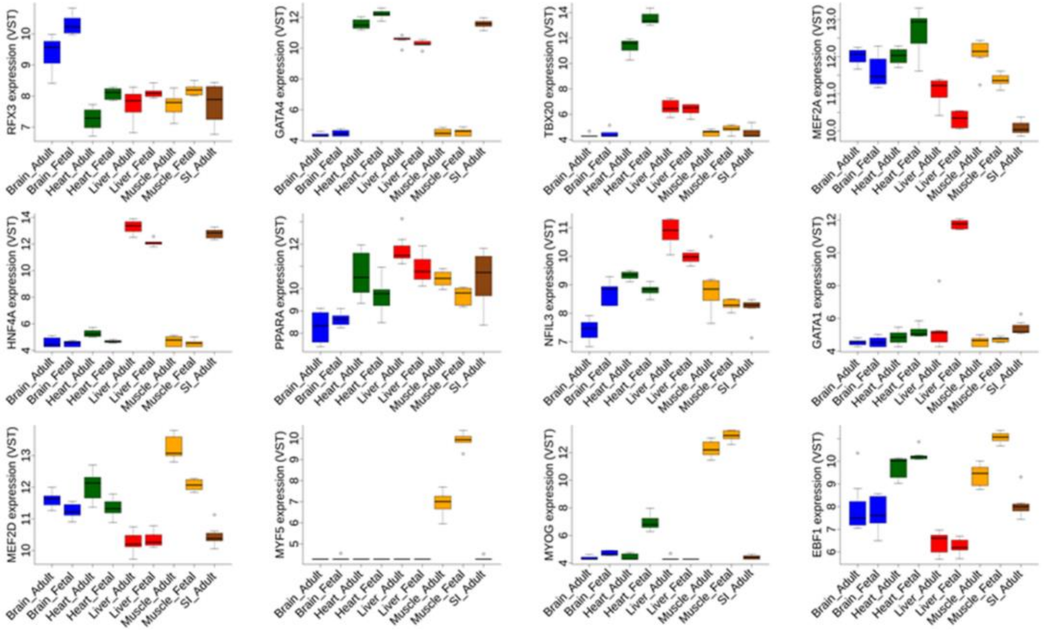


Supplementary Fig. 14: Enrichment of transcription factormotif in developmental stage specific peaks. (A) A table listing a selective transcription factors that were enriched in the developmental stage specific peaks in different tissues, (B) also shown higher gene expression levels in the corresponding tissue-stage..

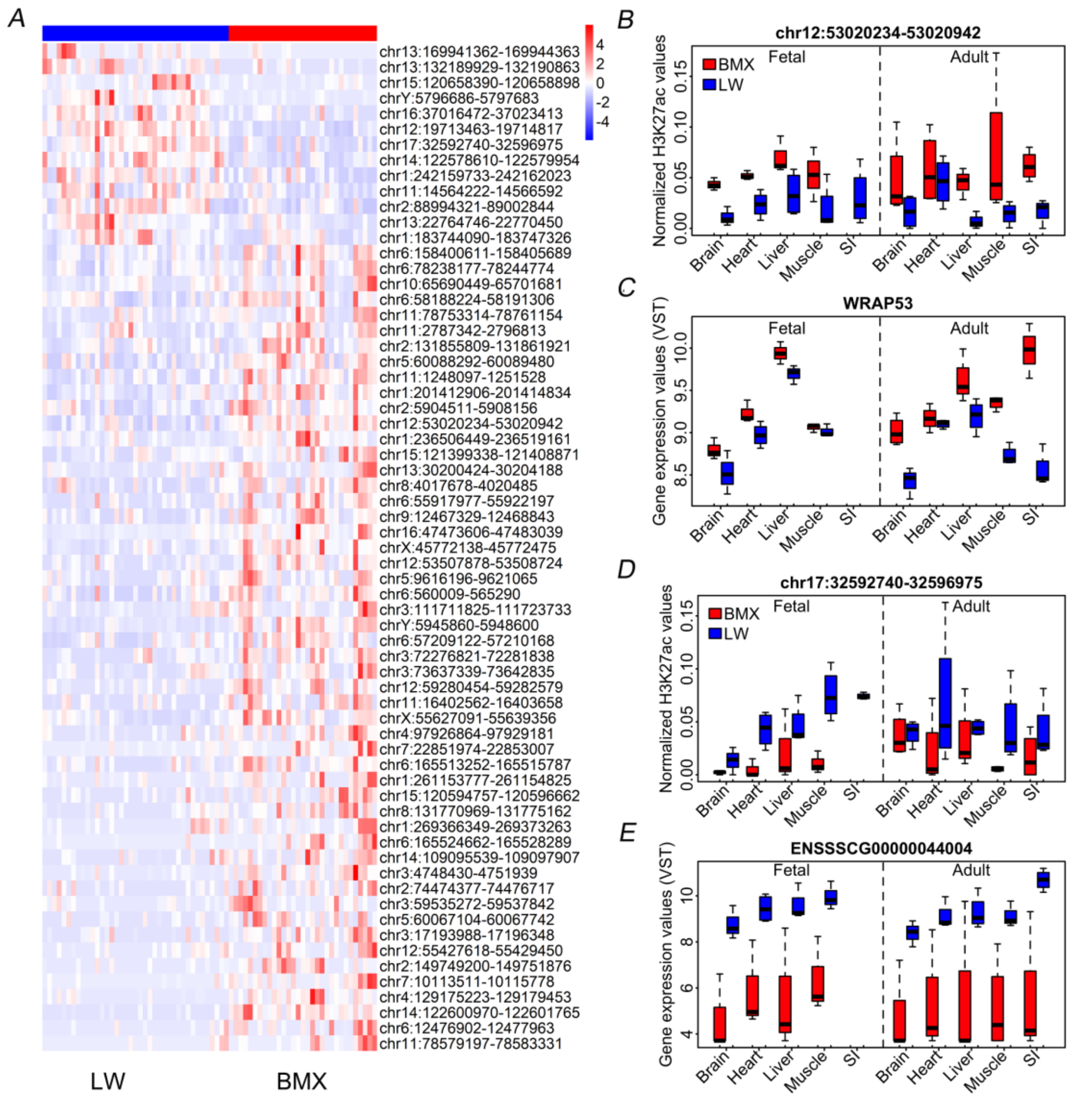


Supplementary Fig. 15**:** H3K27ac peaks showing differential activity between Bama Xiang and Large White pigs across tissue - developmental stages. (A) Heatmap showing the fold change of peak activity in Bama Xiang versus that in Large White across different tissue – developmental stages, the red [26] block in the heatmap represent the results for 52 (13) peaks shown higher (lower) activity in Bama Xiang pigs than those in Large White peaks. (B) Barplot showing greater activity of peak at chr12:53,020,234-53,020,942 in Bama Xiang than in Large White pigs across different tissue – developmental stages. (C) Barplot showing greater expressions of WRAP53 (target gene of the peak at chr12:53,020,234-53,020,942) in Bama

Xiang than in Large White pig across different tissue – developmental stages. (D) Barplot showing lower activity of peak at chr17:32592740-32596975 in Bama Xiang and Large White across different tissue – developmental stages. (E) Barplot showing lower expressions of ENSSSCG00000044004 (target gene of the peak at chr17:32592740-32596975) in Bama Xiang than those in Large White across different tissue – developmental stages.

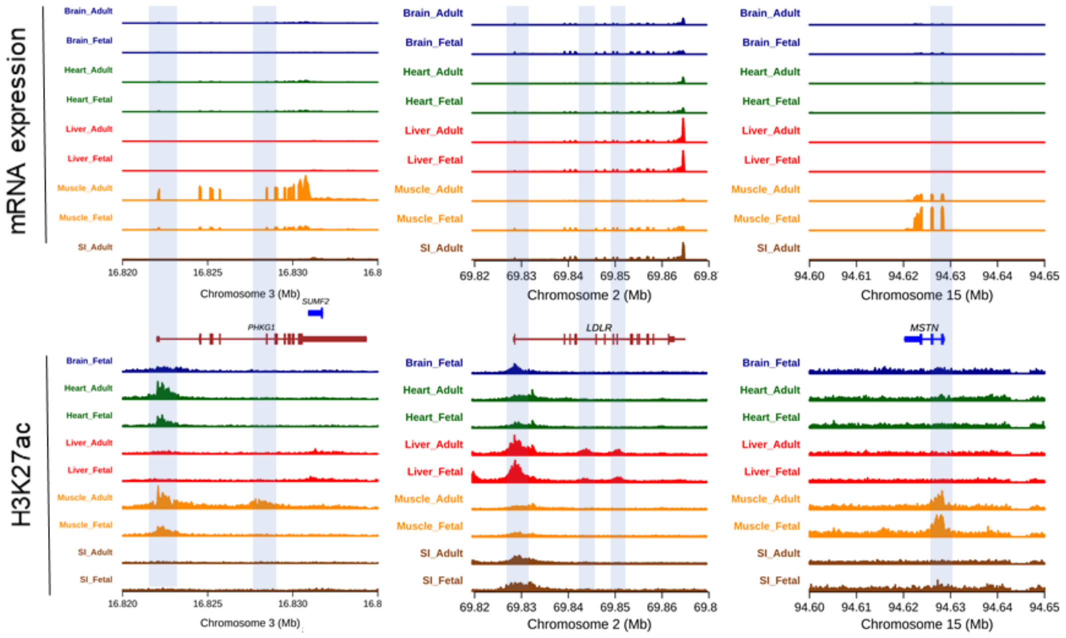


Supplementary Fig. 16: The tracks of gene expression and H3K27ac activity three PHKG1, LDLR and MSTN gene regions across the tissue – developmental stages. The transparent blue bars highlighted the H3K27ac peaks that have potential roles in regulating the corresponding genes.

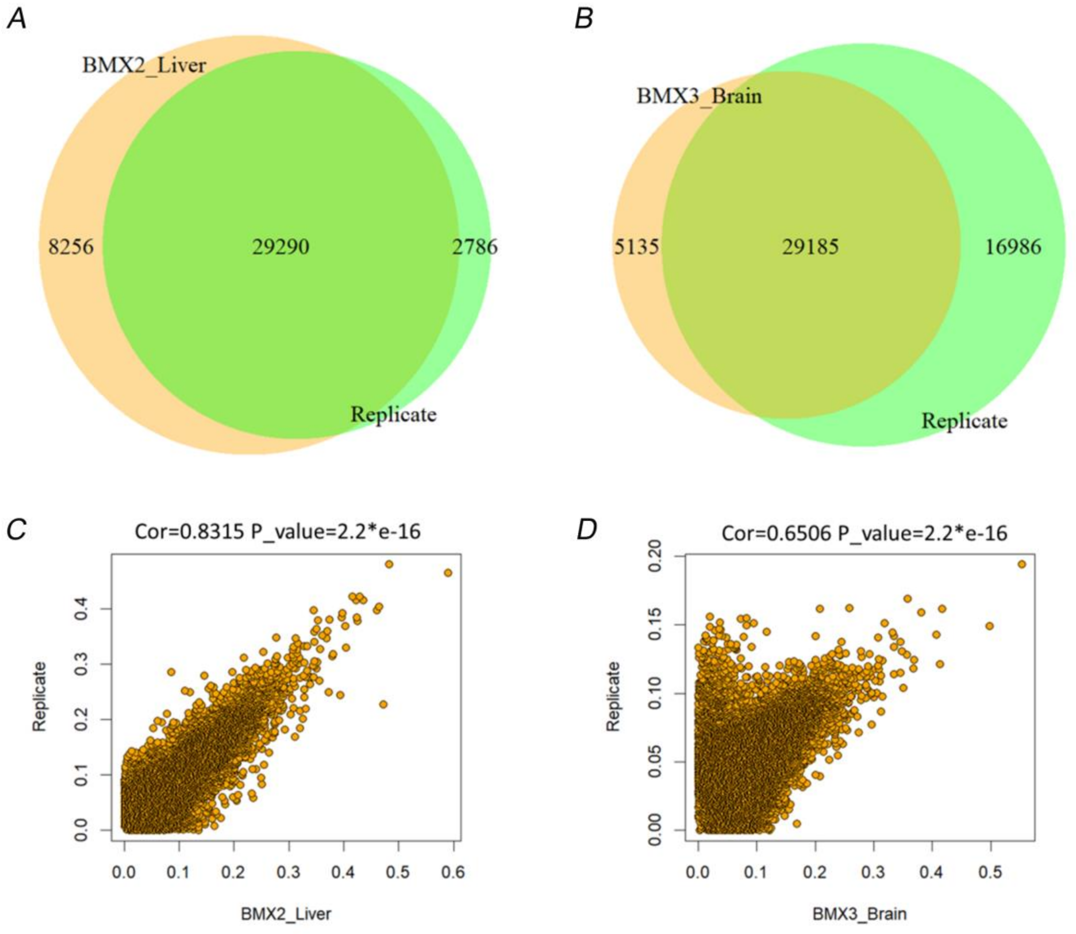


Supplementary Fig. 17: Assessing the data quality using technical replications. The activity of peaks from an adult Bama Xiang liver and an adult brain sample out of the 70 samples were inferred through H3K27ac target ChIP-Seq and following peak calling and quantification pipeline. A-B) Venn plots showing the overlap of peaks between the two technical replicates and their matching samples among the 70 samples, respectively. C-D) Scatter plots showing the correlation of peak activity between the two technical replicates and their matching samples among the 70 samples, respectively.
